# Nuclear TSC2 drives miR-514b-3p transcription to regulate PI3K-AKT-MTOR signalling: Implications for OSCC pathogenesis

**DOI:** 10.64898/2026.03.10.710871

**Authors:** Sakshi Gupta, Naresh Mahajan, Mukund Kumar, Arun Kumar

## Abstract

The PI3K-AKT-MTOR signalling pathway is pivotal in regulating cell survival, proliferation, and growth. TSC2 or tuberin acts as a key negative regulator of this pathway by forming a cytoplasmic complex with TSC1 and TBC1D7. While the cytoplasmic role of TSC2 is well established, emerging evidence suggests its nuclear functions. Previously, we have identified TSC2 as a transcriptional repressor of the *EREG* gene. Building on this foundation, the present study has investigated the transcriptional role of TSC2 in miRNA gene regulation. A genome-wide miRNA microarray profiling of TSC2-depleted SCC131 cells identified 19 upregulated and 24 downregulated miRNAs. Of these, miR-514b-3p emerged as one of the most upregulated miRNAs. The RT-qPCR analysis showed that TSC2 knockdown results in a robust miR-514b-3p upregulation, whereas TSC2 overexpression suppresses its expression in SCC131 cells. Moreover, TSC2 negatively regulates the *MIR514B* promoter activity in an NLS-dependent manner. Our ChIP analysis confirmed the direct binding of TSC2 to the *MIR514B* promoter, establishing miR-514b-3p as a transcriptional target of TSC2. We further identified *TSPAN9* as a direct downstream target of miR-514b-3p. TSC2 positively regulates *TSPAN9* levels by repressing miR-514b-3p, thereby establishing a novel TSC2/miR-514b-3p/TSPAN9 regulatory axis. We further uncovered a crosstalk between TSC2/miR-514b-3p/TSPAN9 axis and the canonical PI3K-AKT-MTOR signalling pathway, where miR-514b-3p positively and *TSPAN9* negatively regulates this pathway. Interestingly, AKT functions as an upstream regulator of this axis by modulating the nuclear localization of TSC2. Collectively, this study provides new insights into the non-canonical, nucleus-dependent transcriptional functions of TSC2, thus expanding its role beyond cytoplasmic signalling regulation and underscoring its significance in the cellular signalling networks.

**Graphical abstract:** **The diagrammatic representation of the TSC2/miR-514b-3p/TSPAN9 axis and its interaction with the canonical PI3K-AKT-MTOR pathway.**

*Abbreviations*: PI3K, Phosphatidylinositol-3 kinase; TSC1, Tuberous sclerosis complex subunit 1; TSC2, Tuberous sclerosis complex subunit 2; TBC1D7, Tre2-Bub2-CDC16 domain family member 7; RHEB, Ras homolog enriched in brain; GTP, Guanosine tri phosphate; GDP, Guanosine di phosphate; MTORC1, Mechanistic target of rapamycin kinase complex 1; S6K1, p70 ribosomal protein S6 kinase; 4E-BP-1, eIF4E-binding protein 1; and TSPAN9, Tetraspanin 9. This figure was created in BioRender (https://app.biorender.com/).

## Introduction

Communication and coordination among different cellular populations are necessary for maintaining homeostasis [1]. In this context, signal transduction pathways act as a molecular language to regulate communication across neighbouring as well as distant cell populations [1]. The PI3K-AKT-MTOR axis, also referred to as the cell survival pathway, is a multi-step signal transduction cascade that regulates diverse cellular processes, including cell growth and proliferation, 5’-cap dependent translation, metabolism, ribosome biogenesis, and apoptosis [2,3]. TSC2 (Tuberous sclerosis complex subunit 2) or tuberin forms a heteromeric cytoplasmic complex with TSC1 and TBC1D7 and acts as a critical cell growth suppressor and a major inhibitor of the PI3K-AKT-MTOR signalling pathway [4,5]. The loss of *TSC2* results in MTORC1 hyperactivation and aberrant cellular growth, due to which *TSC2* is also characterized as a classical tumour suppressor gene [4]. The PI3K-AKT-MTOR pathway occurs outside the nucleus, primarily in the cytoplasm and lysosomal membrane [2]. However several pathway-associated signalling molecules, including TSC2, are also known to translocate into the nucleus, where they contribute to non-canonical nuclear functions [3,6].

The *TSC2* gene is located on chromosome 16p13.3 and codes for an 1,807 aa (amino acids) long protein of ∼200 kDa (https://www.ncbi.nlm.nih.gov/) [4]. In addition to its presence in the cytoplasm and nucleus, TSC2 also resides in mitochondria, lysosome, and the Golgi apparatus, although its compartment-specific functions are not yet clear [3,7–9]. TSC2 is composed of a leucine zipper domain (LZ: aa 75-107), two coiled-coil domains (CC1: aa 346-371 and CC2: 1008-1021), two transcription activation domains (TAD1: aa 1163-1259 and TAD2: aa 1690-1743), a GTPase activating protein homology domain (GAP: aa 1517-1674), and a nuclear localization signal sequence (NLS: aa 1743-1755) [3]. The mutations in *TSC1* and *TSC2* genes cause tuberous sclerosis complex (TSC) disease, an autosomal dominant genetic disorder characterized by the formation of benign tumours (hamartomas) in several organs, including the brain, heart, skin, kidneys, and lungs [10]. Mutations and downregulation of *TSC1* and *TSC2* have also been observed in several cancers, leading to hyperactivation of the MTORC1 signalling and thereby aiding in cancer progression [4]. TSC2 functions as a GAP (GTPase-activating protein) for the small GTPase, RHEB, and converts RHEB to its inactive GDP-bound form, rendering the MTORC1 inactive [5,11]. Apart from its canonical role, TSC2 also exerts cell growth-suppressive effects via MTORC1-independent signalling mechanisms [12]. Although TSC2 is canonically part of the TSC complex, it can carry out non-canonical functions on its own, independent of the complex [12].

TSC2 has also been found in the nucleus of many cell types, including HeLa, HEK293, Rat-1, and NIH3T3, suggesting its ubiquitous nuclear localization [6]. It utilizes a 13 aa-long NLS sequence near its C-terminus (RLRHIKRLRQRIR: aa 1743-1755) for its nuclear localization [13]. Clements et al. [8] have shown that the deletion of aa positions 1351-1807 in TSC2 reduces its nuclear entry in HASM (human airway smooth muscle) cells [8]. Moreover, TSC2 nuclear localization is a phosphorylation-dependent process [14]. The phosphorylation of TSC2 at Ser939 and Thr1462 by AKT inhibits TSC2 nuclear entry and retains it in the cytoplasm [14]. In the nucleus, TSC2 acts as a nuclear receptor co-regulator for the steroid receptor family [15,16]. TSC2 enhances transcription driven by the VDR (vitamin D receptor) and PPAR (peroxisome proliferator-activated receptor), while represses transcriptional activity mediated by the glucocorticoid receptor and ERα (estrogen receptor alpha) [13,15]. In 2014, our research group has found the first evidence of a direct nuclear function of TSC2 as a transcription factor [17]. This study has demonstrated that the nuclear TSC2 binds directly to the promoter of the oncogene *EREG* (Epiregulin) and represses its expression [17]. Interesting, *EREG* has also been found to be upregulated in TSC2 knock out cells from TSC patients [18]. It is a well-accepted premise that *EREG* is unlikely to be the only gene transcriptionally regulated by TSC2. Given the complex role of TSC2 in cellular signalling, it is highly plausible that TSC2 modulates the transcription of multiple other protein-coding as well as non-coding genes. The present study has therefore explored the TSC2-mediated regulation of miRNA (microRNA) genes and the mechanisms underlying its regulation. Our genome-wide miRNA microarray profiling of small RNA enriched samples from TSC2 depleted cells from an OSCC (oral squamous cell carcinoma) cell line SCC131 has revealed several differentially regulated miRNAs. We have chosen one of the most upregulated miRNAs, miR-514b-3p, and have investigated the mechanism of its transcriptional regulation by TSC2 in detail. Since miRNAs exert their effects by binding to target genes, we have also identified its target gene *TSPAN9* and have investigated the role of the TSC2/miR-514b-3p/TSPAN9 axis in the OSCC pathogenesis.

## Results

### Identification of TSC2 as a negative regulator of miR-514b-3p expression

To identify the TSC2-mediated differentially regulated miRNAs, we transiently transfected SCC131 cells with Mock and TSC2 siRNAs pools separately. We then confirmed the TSC2 knockdown at both the mRNA and protein levels (Fig. 1A). As expected, the TSC1 levels remained comparable in both Mock and TSC2 siRNAs transfected cells (Fig. 1A). As *EREG* is a well-established transcriptional target of TSC2, we assessed its expression following TSC2 knockdown. Consistent with the previous report [17], TSC2 silencing resulted in a significant upregulation of *EREG* mRNA levels in TSC2 siRNAs-transfected cells compared to those transfected with Mock siRNAs (Fig. S1). For genome-wide miRNA profiling, small RNA-enriched fractions isolated from both Mock and TSC2 siRNAs-transfected cells were subjected to the miRNA microarray analysis, using the SurePrint G3 8×60K Human miRNA microarray chips with 2,549 mature miRNA probes (Agilent Technologies, Santa Clara, CA, USA). The results showed a total of 43 differentially regulated miRNAs (Fig. 1B, Table S1). Of these, 19 miRNAs were significantly upregulated, and 24 miRNAs were significantly downregulated in cells transfected with TSC2 siRNAs as compared to those transfected with Mock siRNAs (Fig. 1B, Table S1). A heat map representing the differentially regulated miRNAs is shown in Fig. 1B.

**Fig. 1.**
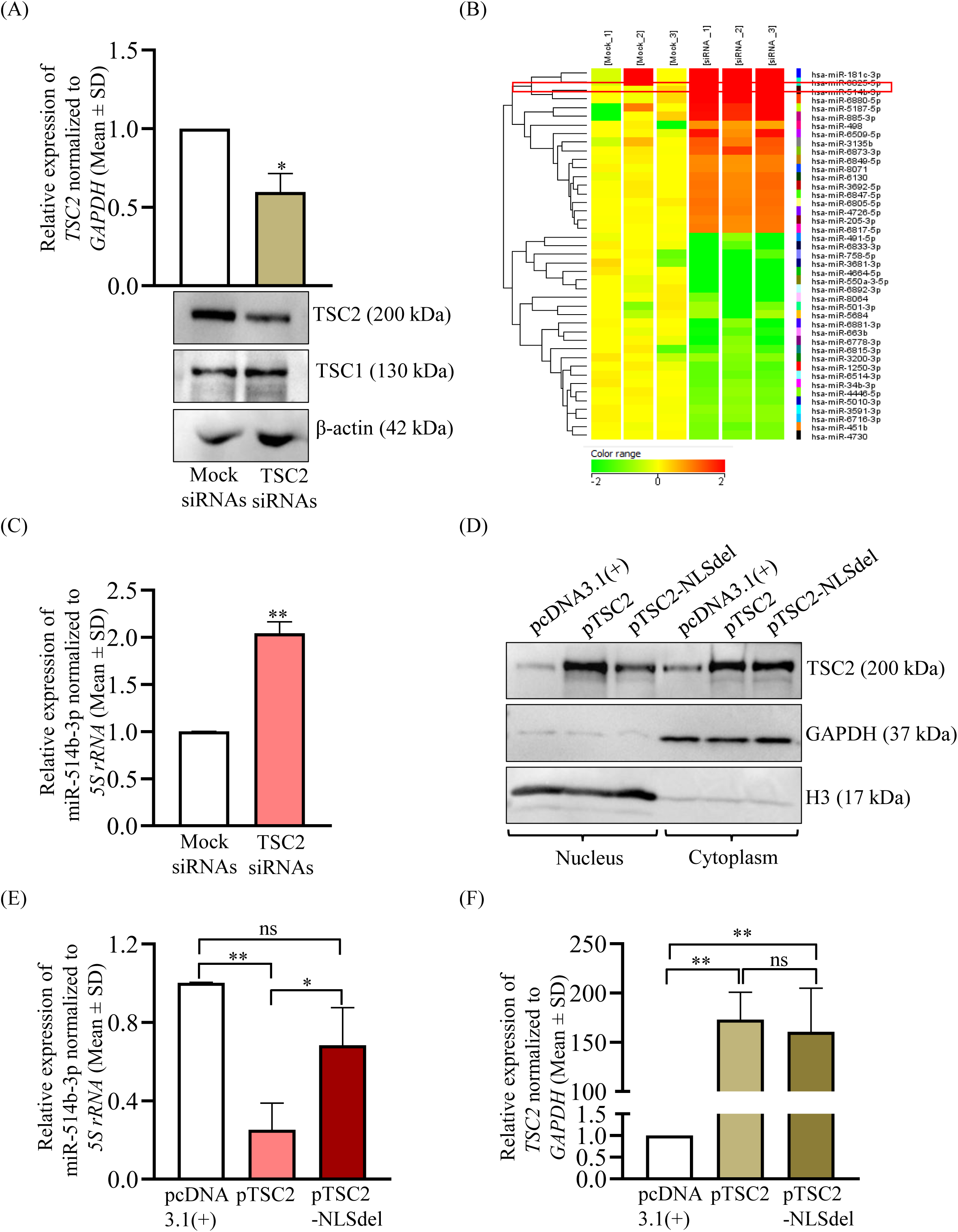
TSC2 negatively regulates miR-514b-3p expression in an NLS-dependent manner. (A) Both the transcript and protein levels of TSC2 were downregulated in cells transfected with TSC2 siRNAs as compared to those transfected with Mock siRNAs. As expected, TSC1 levels remained unchanged in both samples. β-actin was used as a loading control (N=3). (B) A heatmap depicting the upregulated and downregulated miRNAs identified by the miRNA microarray analysis of TSC2 siRNAs and Mock siRNAs-transfected SCC131 cells. Mock_1, Mock_2 and Mock-3 are replicates of small RNAs isolated from cells transfected with Mock siRNAs, whereas siRNA_1, siRNA_2 and siRNA_3 are replicates of small RNAs isolated from cells transfected with TSC2 siRNAs. miR-514b-3p is highlighted in a red box. (C) The upregulation of miR-514b-3p in TSC2 siRNAs-transfected SCC131 cells as compared to those transfected with Mock siRNAs (N=3). (D) Nuclear-cytoplasmic fractionation showing the reduction of nuclear TSC2 in pTSC2-NLSdel-transfected cells as compared to those transfected with pTSC2. GAPDH and H3 were used as cytoplasmic and nuclear markers respectively. (E) The downregulation of miR-514b-3p expression in pTSC2-transfected cells as compared to those transfected with pcDNA3.1(+). Note, the upregulation of miR-514b-3p in pTSC2-NLSdel-transfected cells as compared to those transfected with pTSC2 (N=3). (F) TSC2 overexpression in cells transfected with pTSC2 or pTSC2-NLSdel as compared to those transfected with pcDNA3.1(+) (N=3). P-values were calculated with Unpaired student’s t-test with Welch correction for comparing two data sets and one-way ANOVA, followed by Tukey’s multiple comparison test for multiple comparisons.

Of the 19 upregulated miRNAs, hsa-miR-514b-3p (miR-514b-3p) emerged as one of the most robustly upregulated candidates, exhibiting ∼28.5-fold increase in expression (mean log2 fold change of 4.91) in TSC2-silenced cells as compared to those transfected with the Mock siRNAs (Fig. 1B, Table S1). Owing to its strong upregulation following TSC2 knockdown and the lack of prior information regarding its transcriptional regulation, miR-514b-3p was selected as a potential target of TSC2 and subjected to further investigation. The upregulation of miR-514b-3p post TSC2 knockdown was validated by RT-qPCR and RT-PCR using RT6-miR-514b-3p and shortmiR-514b-3p primers (Figs. 1C, S2, and Table S2). The results showed a significant upregulation in miR-514b-3p expression in TSC2 siRNAs-transfected cells as compared to those transfected with the Mock siRNAs, validating the miRNA microarray data (Figs. 1C and S2). These results suggested that TSC2 negatively regulates the expression of miR-514b-3p.

**Fig. 2.**
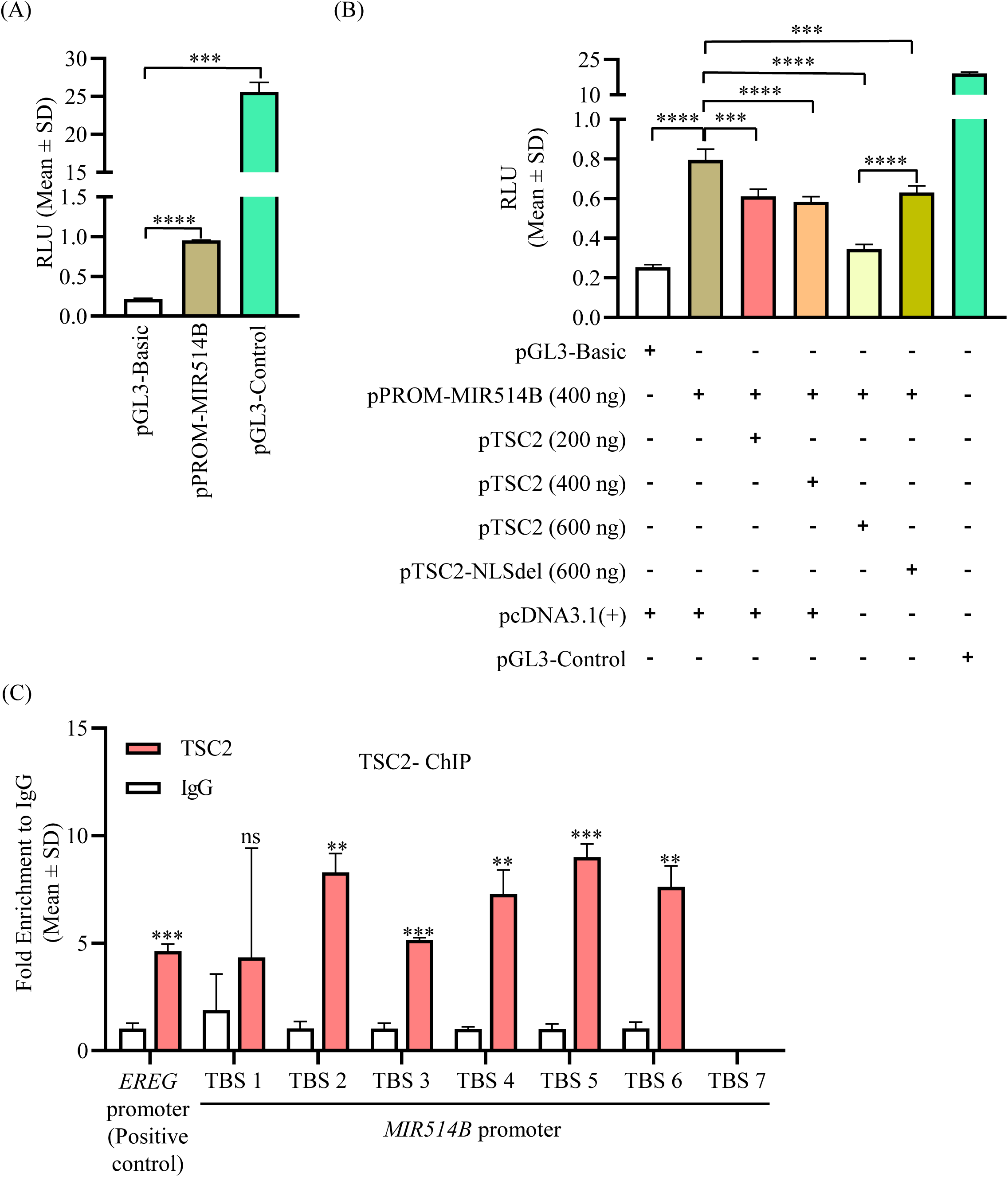
TSC2 acts as a transcriptional repressor of miR-514b-3p. (A) A significant promoter activity was observed in SCC131 cells transfected with pPROM-MIR514B as compared to those transfected with the pGL3-Basic vector. pGL3-Basic served as a negative control and pGL3-Control served as a positive control (N=3). (B) The SCC131 cells with the *MIR514B* promoter construct (pPROM-MIR514B) showed a significantly reduced luciferase activity with an increasing dosage of pTSC2 compared to those co-transfected with pPROM-MIR514B and the pcDNA3.1(+). In addition, the cells with pPROM-MIR514B and pTSC2-NLSdel showed a significantly increased luciferase activity as compared to those co-transfected with pPROM-MIR514B and pTSC2 (N=3). (C) TSC2 shows binding to the five out of seven TBSs (i.e., TBS2, TBS3, TBS4, TBS5, and TBS6) in the *MIR514B* promoter. There was no binding of TSC2 to TBS1, and no amplification was achieved for TBS7. The binding of TSC2 to the *EREG* promoter served as a positive control. All ChIP experiments were plotted as TSC2 fold enrichment in comparison to the IgG control (N=3). P-values were calculated with Unpaired student’s t-test with Welch correction for comparing two data sets and one-way ANOVA, followed by Tukey’s multiple comparison test for multiple comparisons. *Abbreviations*: RLU, Relative light units; TBS, TSC2 binding site; and ChIP, Chromatin immunoprecipitation.

### TSC2 negatively regulates miR-514b-3p expression in an NLS-dependent manner

To determine whether TSC2-mediated repression of miR-514b-3p is dependent on its nuclear localization, we analyzed the miR-514b-3p expression following ectopic expression of wild-type *TSC2* ORF (pTSC2) or an NLS-deleted *TSC2* ORF (pTSC2-NLSdel) in SCC131 cells. Previous studies have demonstrated that TSC2 enters the nucleus via utilizing an NLS sequence near its C-terminus, and deletion of this sequence restricts its nuclear entry [13]. We first validated the NLS-dependent subcellular localization of TSC2 by performing nuclear-cytoplasmic fractionation of SCC131 cells transfected with the vector pcDNA3.1(+), pTSC2 harbouring a full-length *TSC2* ORF, or pTSC2-NLSdel harbouring the *TSC2* ORF with deleted NLS. Consistent with earlier reports [8,13], we also observed a marked reduction in nuclear TSC2 levels in cells transfected with pTSC2-NLSdel as compared to those transfected with pTSC2 (Fig. 1D). The cytoplasmic TSC2 levels remained comparable in cells transfected with pTSC2 and pTSC2-NLSdel (Fig. 1D). These observations confirmed that the deletion of NLS from *TSC2* ORF effectively impairs the entry of TSC2 to the nucleus without substantially affecting its cytoplasmic localization.

We next examined the levels of miR-514b-3p in SCC131 cells transfected with pcDNA3.1(+), pTSC2, or pTSC2-NLSdel. As expected, we observed a significantly reduced level of miR-514b-3p in pTSC2-transfected cells as compared to those transfected with the pcDNA3.1(+), confirming the negative regulation of miR-514b-3p by TSC2 (Fig. 1E). The RT-PCR data also supported this observation (Fig. S2). Interestingly, the level of miR-514b-3p showed a significant increase in cells transfected with pTSC2-NLSdel as compared to those transfected with pTSC2 (Fig. 1E), suggesting that the presence of TSC2 in the nucleus is required for reducing the level of miR-514b-3p, thereby establishing miR-514b-3p regulation as a function of the nuclear TSC2. The levels of miR-514b-3p remained comparable in cells transfected with the vector and pTSC2-NLSdel (Fig. 1E), further suggesting that the nuclear TSC2 is required for the reduced expression of miR-514b-3p. TSC2 overexpression in cells transfected with pTSC2 and pTSC2-NLSdel was confirmed at the mRNA level (Fig. 1F).

### TSC2 acts as a transcriptional repressor for *MIR514B*

Given that TSC2 is a known transcriptional repressor of *EREG*, we speculated whether TSC2 regulates miR-514b-3p at the transcriptional level in a similar manner. Since the promoter of the *MIR514B* gene has not been previously characterized, we first retrieved the putative promoter fragment of 2,825 bp (−2,575 to + 250 bp relative to the transcription start site; Fig. S3A and S3B) from the DBTSS (https://dbtss.hgc.jp/) and the Promoter 2.0 (https://services.healthtech.dtu.dk/services/Promoter-2.0/) databases, and then functionally validated the cloned fragment pPROM-MIR514B by the dual-luciferase reporter assay in SCC131 cells. The results showed a significant luciferase activity in cells transfected with pPROM-MIR514B as compared to those transfected with the pGL3-Basic vector (Fig. 2A), suggesting that the putative fragment indeed represents the *MIR514B* gene promoter. We then aimed to examine the effect of pTSC2 and pTSC2-NLSdel on the *MIR514B* promoter (pPROM-MIR514B) activity by the dual-luciferase reporter assay. To this end, we transfected SCC131 cells with pPROM-MIR514B alone, co-transfected with pPROM-MIR514B and different quantities of pTSC2 (i.e., 200 ng, 400 ng, and 600 ng), or co-transfected with pPROM-MIR514B and pTSC2-NLSdel. The subsequent results showed concomitant reduction in the luciferase activity in cells co-transfected with pPROM-MIR514B and different quantities of pTSC2 in a dose-dependent manner as compared to those transfected with pPROM-MIR514B only, suggesting that TSC2 negatively regulates the *MIR514B* promoter activity (Fig. 2B). Furthermore, the luciferase activity significantly increased in cells co-transfected with pPROM-MIR514B (400 ng) and pTSC2-NLSdel (600 ng) as compared to those co-transfected with pPROM-MIR514B (400 ng) and pTSC2 (600 ng), suggesting that TSC2 negatively regulates the *MIR514B* promoter activity in a nucleus-dependent manner (Fig. 2B). Similar results were observed in cells from another OSCC cell line, SCC084 (Fig. S4).

To determine whether TSC2 acts as a transcriptional repressor of *MIR514B* by directly binding to its promoter, we performed the ChIP-qPCR analysis. Previous research from our group had identified a putative TBS (TSC2 binding site) 5’ GCCTTG 3’ in the promoters of several TSC2-regulated genes [17]. We therefore looked for this putative TBS in the characterized *MIR514B* promoter near the TSS and found seven partial TBSs: TBS1-TBS7 (Fig. S5A and S5B). We designed separate primers across all seven potential TBSs and amplified them individually by ChIP-qPCR, using DNA isolated from TSC2 and IgG pull-down samples (Table S3). The ChIP-qPCR analysis showed strong TSC2 enrichments at five out of seven TBSs in the *MIR514B* promoter (viz., TBS2, TBS3, TBS4, TBS5, and TBS6) in the TSC2 pull-down sample as compared to the IgG pull-down sample, confirming the direct binding of TSC2 to the *MIR514B* promoter (Fig. 2C). Further, TBS1 did not show any TSC2 enrichment, whereas TBS7 did not show any amplification in ChIP-qPCR (Fig. 2C). As expected, a strong TSC2 enrichment was observed at the *EREG* promoter in the TSC2 pull-down sample as compared to the IgG pull-down sample (Fig. 2C). A H3-ChIP was also performed which served as an internal positive control (Fig. S6). The TSC2 enrichment at multiple sites in the *MIR514B* promoter confirmed the direct binding of TSC2 to the *MIR514B* promoter. The above observations suggested that TSC2 acts as a transcriptional repressor by binding directly to the *MIR514B* promoter and negatively regulating the expression of miR-514b-3p.

### *TSPAN9* is a direct gene target of miR-514b-3p

The miRNAs bind to their cognate target mRNAs via sequence complementarity to regulate their gene expression. To find out the direct gene targets of miR-514b-3p, we employed five miRNA prediction tools and observed several putative target genes (Table S4). Based on the rank of appearance in the prediction tools and a thorough literature survey regarding their known functions, we selected following five putative miR-514b-3p target genes: *TSPAN9* (Tetraspanin 9)*, PTEN* (Phosphatase and Tensin Homolog deleted on chromosome 10)*, GULP1* (GULP PTB Domain Containing Engulfment Adaptor 1)*, SESN3* (Sestrin 3), and *CABLES1* (Cdk5 and Abl Enzyme Substrate 1) (Table S4). To determine if these putative genes are targeted by miR-514b-3p, we examined their transcript levels in an miR-514b-3p overexpression background (Fig. S7), along with a known target gene *RBX* (Ring-box 1) [19]. Only two genes, *TSPAN9* and *GULP1* showed a significant downregulation of their transcript levels in SCC131 cells transfected with pMIR514B as compared to those transfected with the pcDNA3-EGFP (Fig. S7). As expected, the transcript level of *RBX* showed a significant downregulation post miR-514b-3p overexpression (Fig. S7). Since the transcript level of *TSPAN9* was significantly more downregulated than that of *GULP1*, we decided to further analyse the interaction of *TSPAN9* and miR-514b-3p in details. To determine if miR-514b-3p also regulates the TSPAN9 protein levels, we transiently transfected SCC131 cells with pMIR514B and pcDNA3-EGFP separately. The results showed a marked reduction in TSPAN9 protein levels in cells transfected with pMIR514B compared to those transfected with the pcDNA3-EGFP, suggesting that miR-514b-3p negatively regulates the protein level of TSPAN9 (Fig. 3A). To confirm this observation further, we performed a dose-dependent transient transfection of SCC131 cells with different quantities of the pMIR514B construct (viz., 2µg, 4µg, and 6µg) and pcDNA3-EGFP separately and determined the transcript and protein levels of TSPAN9. The results showed a concomitant reduction in TSPAN9 transcript and protein levels with increasing levels of miR-514b-3p, suggesting a dose-dependent negative regulation of TSPAN9 by miR-514b-3p (Fig. 3B).

**Fig. 3.**
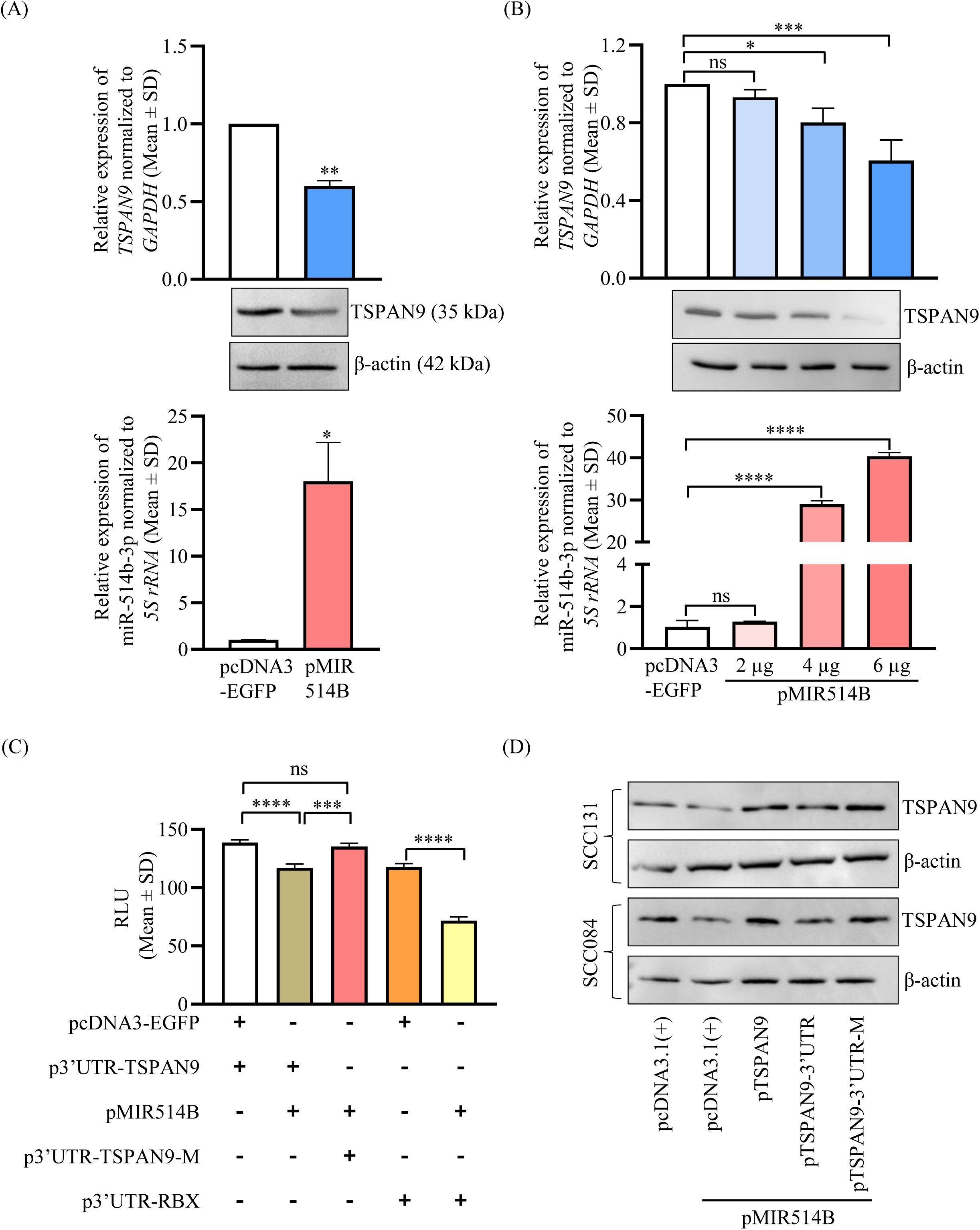
*TSPAN9* is a direct target of miR-514b-3p. (A) The downregulation of TSPAN9 at both the transcript and protein levels in SCC131 cells transfected with pMIR514B as compared to those transfected with pcDNA3-EGFP (upper and middle panels). Note, upregulation of miR-514b-3p in cells transfected with pMIR514B as compared to those transfected with pcDNA3-EGFP (lower panel). β-actin was used as a loading control (N=3). (B) TSPAN9 showed a concomitant reduction at both transcript and protein levels with an increase in miR-514b-3p expression in a dose-dependent manner (upper and middle panels). Note, concomitant upregulation of miR-514b-3p in cells transfected with pMIR514B as compared to those transfected with pcDNA3-EGFP (lower panel). β-actin was used as loading control (N=3). (C) The dual-luciferase reporter assay showing a direct interaction between the TS in the 3’UTR of *TSPAN9* and the SR of miR-514b-3p in SCC131 cells. Note, a significant decrease in the luciferase activity in cells co-transfected with p3’UTR-TSPAN9 and pMIR514B compared to those transfected with p3’UTR-TSPAN9 and pcDNA3-EGFP. As expected, the cells co-transfected with p3’UTR-RBX and pMIR514B showed a significantly reduced luciferase activity compared to those co-transfected with p3’UTR-RBX and pcDNA3-EGFP (N=3). (D) The Western blot analysis showing the levels of TSPAN9 in SCC131 cells transfected with pcDNA3.1(+) only or co-transfected with pMIR514B and pcDNA3.1(+), pMIR514B and pTSPAN9, pMIR514B and pTSPAN9-3’UTR, or pMIR514B and pTSPAN9-3’UTR-M. β-actin was used as loading control. P-values were calculated with Unpaired student’s t-test with Welch correction for comparing two data sets and one-way ANOVA, followed by Tukey’s multiple comparison test for multiple comparisons. *Abbreviations*: RLU, Relative light units

Our bioinformatics analysis predicted a putative TS (target site) for miR-514b-3p in the 3’UTR of *TSPAN9,* spanning nucleotides 3432-3439, which is conserved across species (Fig. S8). We performed a dual-luciferase reporter assay to confirm the direct interaction between the SR (seed region) of miR-514b-3p and the TS in the 3’UTR of *TSPAN9*, using constructs illustrated in Fig. S9A. The p3’UTR-TSPAN9 construct harbours *TSPAN9* 3’UTR in sense orientation with the intact TS, whereas the p3’UTR-TSPAN9-M construct is generated by abrogating the TS for miR-514b-3p in the 3’UTR of *TSPAN9* by site-directed mutagenesis (Fig. S9A). To determine whether miR-514b-3p directly interacts with TS present in the 3’UTR of *TSPAN9*, we co-transfected SCC131 cells separately with p3’UTR-TSPAN9 and pcDNA3-EGFP, p3’UTR-TSPAN9 and pMIR514B, or p3’UTR-TSPAN9-M and pMIR514B and performed a dual-luciferase reporter assay. Compared to cells co-transfected with p3’UTR-TSPAN9 and pcDNA3-EGFP, we observed a significant decrease in the luciferase activity in those co-transfected with p3’UTR-TSPAN9 and pMIR514B, confirming that miR-514b-3p binds directly to the 3’UTR of *TSPAN9* (Fig. 3C). Moreover, cells co-transfected with p3’UTR-TSPAN9-M and pMIR514B showed the luciferase activity comparable to those co-transfected with p3’UTR-TSPAN9 and pcDNA3-EGFP, due to the absence of miR-514b-3p TS in the p3’UTR-TSPAN9-M construct (Fig. 3C). These observations confirmed that the miR-514b-3p binds directly to the TS present in the *TSPAN9* 3’UTR in a sequence-specific manner. We also observed a direct interaction between miR-514b-3p and *RBX* 3’UTR, which served as a positive control for the assay (Fig. 3C).

### miR-514b-3p negatively regulates TSPAN9 protein levels by interacting with its 3’UTR

We have so far demonstrated that miR-514b-3p negatively regulates TSPAN9 expression and exhibits a direct binding to the 3’UTR of *TSPAN9* in a sequence-specific manner. To unequivocally show that the binding between miR-514b-3p and *TSPAN9* 3’UTR downregulates TSPAN9 expression, we performed the Western blot analysis with the following constructs: pTSPAN9 (harbours *TSPAN9* ORF), pTSPAN9-3’UTR (harbours *TSPAN9* ORF and its 3’UTR with a functional TS for miR-514b-3p), and pTSPAN9-3’UTR-M (harbours *TSPAN9* ORF and its 3’UTR with abrogated TS for miR-514b-3p) (Fig. S9B). We transiently transfected SCC131 and SCC084 cells with pcDNA3.1(+) only or co-transfected with pcDNA3.1(+) and pMIR514B, pTSPAN9 and pMIR514B, pTSPAN9-3’UTR and pMIR514B, or pTSPAN9-3’UTR-M and pMIR514B and subsequently determined the TSPAN9 protein levels. The results showed that, as compared to cells transfected with pcDNA3.1(+) only, cells co-transfected with pcDNA3.1(+) and pMIR514B showed a decreased level of TSPAN9, due to targeting of the endogenous *TSPAN9* by miR-514b-3p (Fig. 3D). Further, TSPAN9 levels were increased in cells co-transfected with pTSPAN9 and pMIR514B as compared to those co-transfected with pcDNA3.1(+) and pMIR514B, due to the TSPAN9 overexpression by pTSPAN9 (Fig. 3D). Furthermore, the cells co-transfected with pTSPAN9-3’UTR and pMIR514B showed a marked reduction in the TSPAN9 levels as compared to those transfected with pTSPAN9 and pMIR514B, due to the presence of a functional TS in the pTSPAN9-3’UTR, resulting in the interaction between the 3’UTR of *TSPAN9* and miR-514b-3p (Fig. 3D). Moreover, the TSPAN9 levels in cells co-transfected with pTSPAN9-3’UTR-M and pMIR514B were comparable to those transfected with pTSPAN9 and pMIR514B, due to the abrogation of TS in pTSPAN9-3’UTR-M (Fig. 3D). These observations confirmed that the negative regulation of *TSPAN9* by miR-514b-3p is dependent on the presence or absence of its 3’UTR.

### TSC2 positively regulates *TSPAN9* expression via the TSC2/miR-514b-3p/TSPAN9 axis

Since TSC2 negatively regulates miR-514b-3p, which in turn negatively regulates *TSPAN9*, it is reasonable to assume that TSC2 might positively regulate *TSPAN9* expression. To this end, we performed TSC2 knockdown and TSC2 overexpression in SCC131 cells. As expected, we observed a sharp reduction in the TSPAN9 expression at both transcript and protein levels in cells transfected with TSC2 siRNAs compared to those transfected with the Mock siRNAs (Fig. 4A). Furthermore, TSPAN9 was upregulated in cells transfected with pTSC2 compared to those transfected with pcDNA3.1(+), suggesting TSC2-mediated positive regulation of TSPAN9 at both the transcript and protein levels (Fig. 4B). Moreover, the TSPAN9 expression in pTSC2-NLSdel-transfected SCC131 cells was comparable to those transfected with the vector pcDNA3.1(+), suggesting TSC2-mediated NLS-dependent regulation of TSPAN9 expression (Fig. 4B). These observations suggested the existence of a nucleus-dependent TSC2/miR-514b-3p/TSPAN9 axis.

**Fig. 4.**
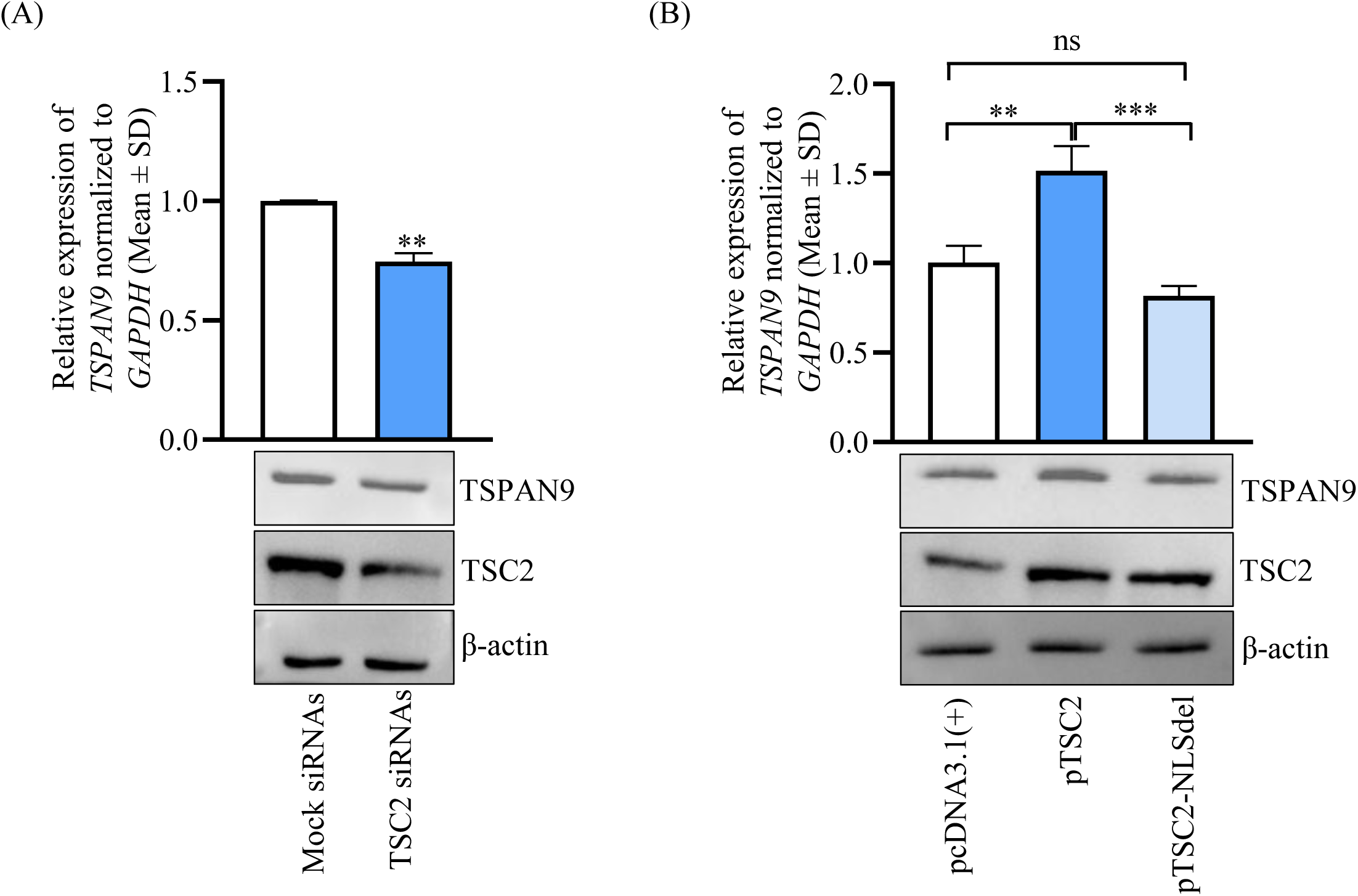
TSC2 positively regulates the *TSPAN9* expression in an NLS-dependent manner. A) *TSPAN9* showed downregulation in TSC2 siRNAs-transfected SCC131 cells compared to the Mock siRNAs-transfected cells at both transcript and protein levels. Note, TSC2 knockdown was confirmed at protein level (lower panel) (N=3). B) *TSPAN9* showed upregulation in pTSC2-transfected SCC131 cells compared to pcDNA3.1(+)-transfected cells at both transcript and protein levels. Note, *TSPAN9* expression was comparable between pTSC2-NLSdel-transfected cells and pcDNA3.1(+)-transfected cells. TSC2 overexpression was confirmed at the protein level (lower panel) (N=3). P-values were calculated with Unpaired student’s t-test with Welch correction for comparing two data sets and one-way ANOVA, followed by Tukey’s multiple comparison test for multiple comparisons.

### TSC2 and TSPAN9 are anti-proliferative, whereas miR-514b-3p is pro-proliferative in nature

After establishing the TSC2/miR-514b-3p/TSPAN9 axis, we aimed to examine the independent effects of each of these molecules on the proliferation of OSCC cells. We transfected OSCC cells separately with pTSC2, pTSPAN9, and pMIR514B along with appropriate vectors as controls and assessed proliferation using the trypan blue dye exclusion and BrdU cell proliferation assays. The trypan blue dye exclusion assay showed a significantly reduced proliferation of OSCC cells transfected with pTSC2 or pTSPAN9 as compared to those transfected with pcDNA3.1(+), suggesting the anti-proliferative nature of TSC2 and TSPAN9 (Figs. 5A, 5B, S10A, and S10B). Further, we observed a significant increase in the proliferation of OSCC cells transfected with pMIR514B compared to those transfected with pcDNA3-EGFP, indicating the pro-proliferative nature of miR-514b-3p (Figs. 5C and S10C). The BrdU cell proliferation assay also showed a significant reduction in absorbance in OSCC cells transfected with pTSC2 or pTSPAN9 as compared to those transfected with pcDNA3.1(+), confirming the anti-proliferative nature of TSC2 and TSPAN9 (Figs. 5D, 5E, S10D, and S10E). Further, we observed a significant increase in absorbance in OSCC cells transfected with pMIR514B as compared to those transfected with pcDNA3-EGFP, confirming the pro-proliferative nature of miR-514b-3p (Figs. 5F and S10F).

**Fig. 5.**
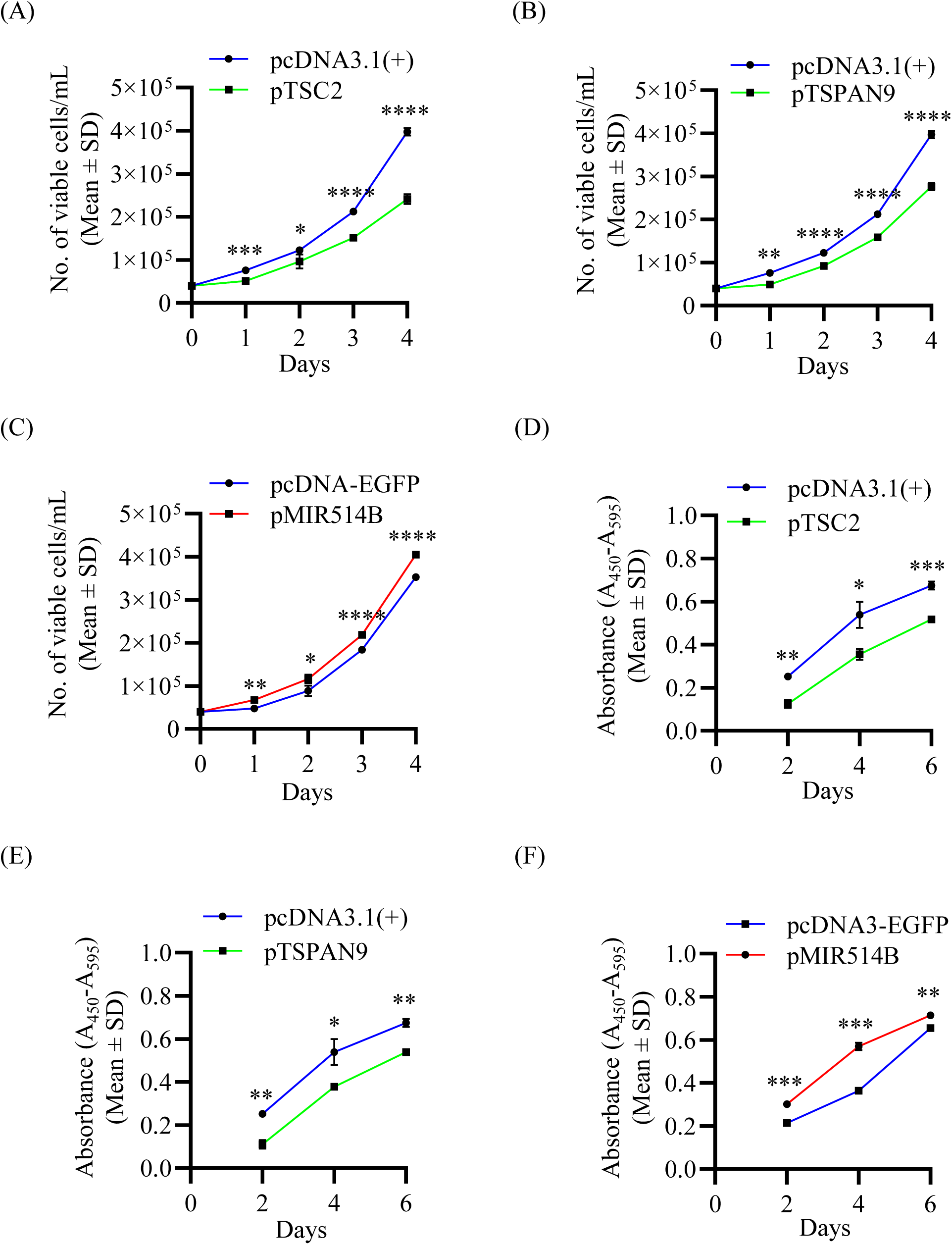
Effects of TSC2, TSPAN9 and miR-514b-3p on proliferation of SCC131 cells. (A-C) The trypan blue dye exclusion assay showing the rate of proliferation of cells transfected with pTSC2, pTSPAN9 or pMIR514B (N=4). (D-F) The BrdU cell proliferation assay showing the rate of proliferation of cells transfected with pTSC2, pTSPAN9 or pMIR514B (N=3). P-values were calculated with Unpaired student’s t-test with Welch correction.

### miR-514b-3p regulates the proliferation, apoptosis, and anchorage-independent growth of OSCC cells, in part, by targeting the 3’UTR of *TSPAN9*

To determine if miR-514b-3p regulates different hallmarks of cancer by targeting the 3’UTR of *TSPAN9*, we transiently transfected SCC131 and SCC084 cells with different constructs in the following combinations: pcDNA3.1(+) only, pcDNA3.1(+) and pMIR514B, pTSPAN9 and pMIR514B, pTSPAN9-3’UTR and pMIR514B, or pTSPAN9-3’UTR-M and pMIR514B and performed assays for different cancer hallmarks.

The trypan blue dye exclusion assay showed that the cells co-transfected with pcDNA3.1(+) and pMIR514B showed an enhanced cell proliferation as compared to those transfected with the pcDNA3.1(+) only, indicating the pro-proliferative nature of miR-514b-3p (Figs. 6A and S11A). The cells co-transfected with pTSPAN9-3’UTR and pMIR514B showed an increased cell proliferation as compared to those transfected with pTSPAN9 and pMIR514B, as well as pTSPAN9-3’UTR-M and pMIR514B, due to the functional TS in pTSPAN9-3’UTR (Figs. 6A and S11A). Moreover, the proliferation of cells co-transfected with pTSPAN9-3’UTR-M and pMIR514B was comparable to those co-transfected with pTSPAN9 and pMIR514B, due to the absence of a functional TS in pTSPAN9-3’UTR-M (Figs. 6A and S11A). Similar results were observed by the BrdU cell proliferation assay in both cell lines (Figs. 6B and S11B). These observations clearly suggest that miR-514b-3p positively regulates the proliferation of OSCC cells, in part, by targeting the 3’UTR of *TSPAN9*.

**Fig. 6.**
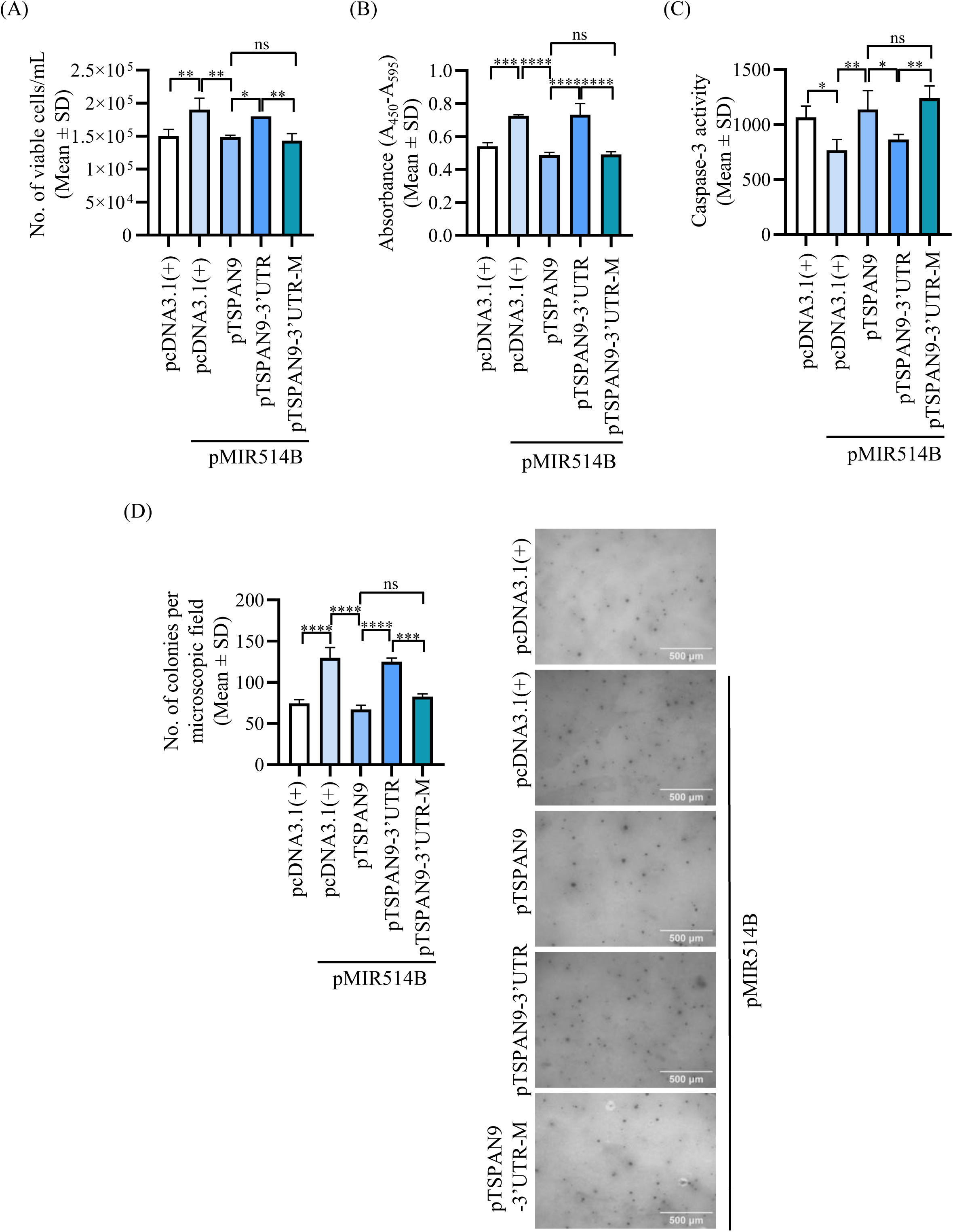
miR-514b-3p regulates proliferation, apoptosis, and anchorage-independent growth of SCC131 cells, in part, by targeting the 3’UTR of *TSPAN9.* (A) Effects of miR-514b-3p and different *TSPAN9* constructs on proliferation of SCC131 cells by the trypan blue dye exclusion assay (N=3) and (B) by the BrdU cell proliferation assay (N=3). (C) Effects of miR-514b-3p and different *TSPAN9* constructs on apoptosis by the Caspase-3 activation assay in SCC131 cells (N=4). (D) Effects of miR-514b-3p and different *TSPAN9* constructs on anchorage-independent growth of SCC131 cells (left panel). Microscopic images of the soft agar colony formation assay are also shown (right panel) (N=3). P-values were calculated by one-way ANOVA, followed by Tukey’s multiple comparison test.

We next examined apoptosis of OSCC cells by the Caspase-3 activation assay. The results demonstrated that the cells co-transfected with pcDNA3.1(+) and pMIR514B showed a reduced Caspase-3 activity as compared to those transfected with only pcDNA3.1(+), indicating the anti-apoptotic nature of miR-514b-3p (Figs. 6C and S11C). The cells co-transfected with pTSPAN9-3’UTR and pMIR514B showed a reduced Caspase-3 activity as compared to those transfected with pTSPAN9 and pMIR514B as well as pTSPAN9-3’UTR-M and pMIR514B, due to a functional TS in pTSPAN9-3’UTR (Figs. 6C and S11C). Moreover, the Caspase-3 activity in cells co-transfected with pTSPAN9-3’UTR-M and pMIR514B was comparable to those co-transfected with pTSPAN9 and pMIR514B, due to the absence of a functional TS in the 3’UTR of *TSPAN9* in pTSPAN9-3’UTR-M (Figs. 6C and S11C). The above observations suggested that miR-514b-3p negatively regulates apoptosis of OSCC cells, in part, by targeting the 3’UTR of *TSPAN9*.

To study the anchorage-independent growth of OSCC cells, we performed a soft agar colony formation assay and observed that cells co-transfected with pcDNA3.1(+) and pMIR514B showed an enhanced colony formation as compared to those transfected with only pcDNA3.1(+), indicating a pro-proliferative nature of miR-514b-3p (Figs. 6D and S11D). The cells co-transfected with pTSPAN9-3’UTR and pMIR514B showed an increased number of colonies as compared to those transfected with pTSPAN9 and pMIR514B, as well as pTSPAN9-3’UTR-M and pMIR514B, due to the functional TS in pTSPAN9-3’UTR (Figs. 6D and S11D). Moreover, the number of colonies in cells co-transfected with pTSPAN9-3’UTR-M and pMIR514B was comparable to those co-transfected with pTSPAN9 and pMIR514B, due to the absence of a functional TS in pTSPAN9-3’UTR-M (Figs. 6D and S11D). Thus, the miR-514b-3p-mediated inhibition of *TSPAN9* leads to an enhanced anchorage-independent growth of OSCC cells due to the pro-proliferative nature of miR-514b-3p, suggesting that miR-514b-3p positively regulates the anchorage-independent growth of OSCC cells, in part, by targeting the 3’UTR of *TSPAN9*.

### The miR-514b-3p/TSPAN9 axis regulates the PI3K-AKT-MTOR pathway via nucleus

TSC2 is a well-established suppressor of the PI3K-AKT-MTOR pathway. As we have demonstrated that the regulation of miR-514b-3p/TSPAN9 is a nuclear function of TSC2, we were then interested to know if the downstream targets of the nuclear TSC2 (viz., miR-514b-3p and TSPAN9) can also regulate the canonical PI3K-AKT-MTOR pathway. To this end, we overexpressed miR-514b-3p and TSPAN9 along with their appropriate vector controls in OSCC cells and analyzed the PI3K-AKT-MTOR pathway readout by checking the levels of phospho-S6K1 (Thr389) (pS6K1) and total S6K1. We also analyzed the levels of crucial mediators such as PI3K (p55), phospho-AKT (Ser473) (pAKT), and total AKT. Following miR-514b-3p overexpression, we observed increased levels of pS6K1, p55, and pAKT in OSCC cells transfected with pMIR514B as compared to those transfected with pcDNA3-EGFP, suggesting that miR-514b-3p positively regulates the PI3K-AKT-MTOR pathway (Fig. 7A). The total S6K1 and total AKT levels remained similar in both samples (Fig. 7A). As expected, TSPAN9 level exhibited a marked decrease in cells transfected with pMIR514B as compared to those transfected with the pcDNA3-EGFP due to miR-514b-3p-mediated TSPAN9 inhibition (Fig. 7A).

**Fig. 7.**
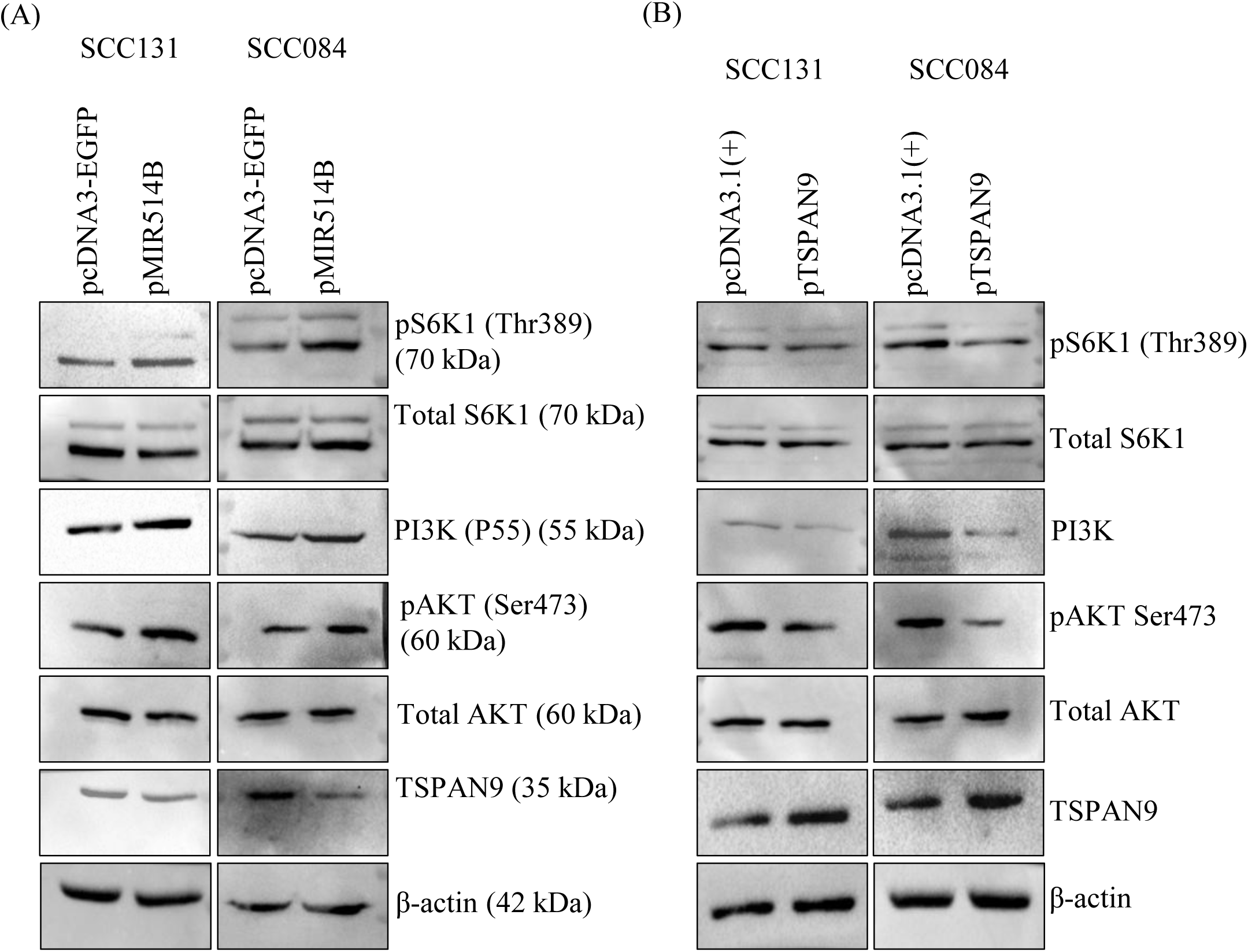
The miR-514b-3p/TSPAN9 axis regulates the PI3K-AKT-MTOR pathway in OSCC cells. (A) The miR-514b-3p overexpression showed increased levels of pS6K1, PI3K (p55), and pAKT in pMIR514B-transfected cells compared to those transfected with pcDNA3-EGFP. (B) The *TSPAN9* overexpression showed decreased levels of pS6K1, PI3K (p55), and pAKT in pTSPAN9-transfected cells compared to those transfected with pcDNA3.1(+). β-actin was used as loading control.

Furthermore, we observed decreased levels of pS6K1, p55, and pAKT in cells transfected with pTSPAN9 as compared to those transfected with pcDNA3.1(+), suggesting that TSPAN9 negatively regulates the PI3K-AKT-MTOR pathway (Fig. 7B). The total S6K1 and total AKT levels remained similar in both samples (Fig. 7B). It is important to note that the activation of the PI3K-AKT-MTOR pathway is widely known to enhance cell proliferation. Our observations thus align well with the pro-proliferative nature of miR-514b-3p and the anti-proliferative nature of *TSPAN9* (Figs. 5 and S10). These observations therefore suggested that miR-514b-3p is a positive regulator, whereas TSPAN9 is a negative regulator of the PI3K-AKT-MTOR pathway.

### AKT modulates the TSC2-miR-514b-3p-TSPAN9 axis via TSC2 nuclear localization

The phosphorylation of TSC2 by AKT at Ser939 and Thr1462 leads to its inactivation and proteasomal degradation [19]. Interestingly, AKT phosphorylation is also crucial for TSC2 nuclear-cytoplasmic shuttle dynamics, where the AKT-mediated inactivating phosphorylation of TSC2 reduces its nuclear entry [19]. To understand the role of AKT in TSC2 nuclear localization in our model system and its effect on the TSC2/miR-514b-3p/TSPAN9 axis, we have used AKT inhibitor VIII (AKTi) to inactivate AKT activity in SCC131 and SCC084 cells. Our hypothesis was based on the premise that the AKT inactivation should drive more TSC2 to the nucleus, which might further inhibit miR-514b-3p expression transcriptionally, and thus enhance TSPAN9 levels.

To confirm the AKT-mediated nuclear localization of TSC2, we performed nuclear-cytoplasmic fractionation of AKTi and DMSO (vehicle control)-treated OSCC cells, followed by the Western blot analysis. The results showed increased levels of nuclear TSC2 in AKTi-treated cells as compared to those treated with the DMSO, whereas the cytoplasmic pool of TSC2 remained unchanged in both treatments (Fig. 8; upper panels).

**Fig. 8.**
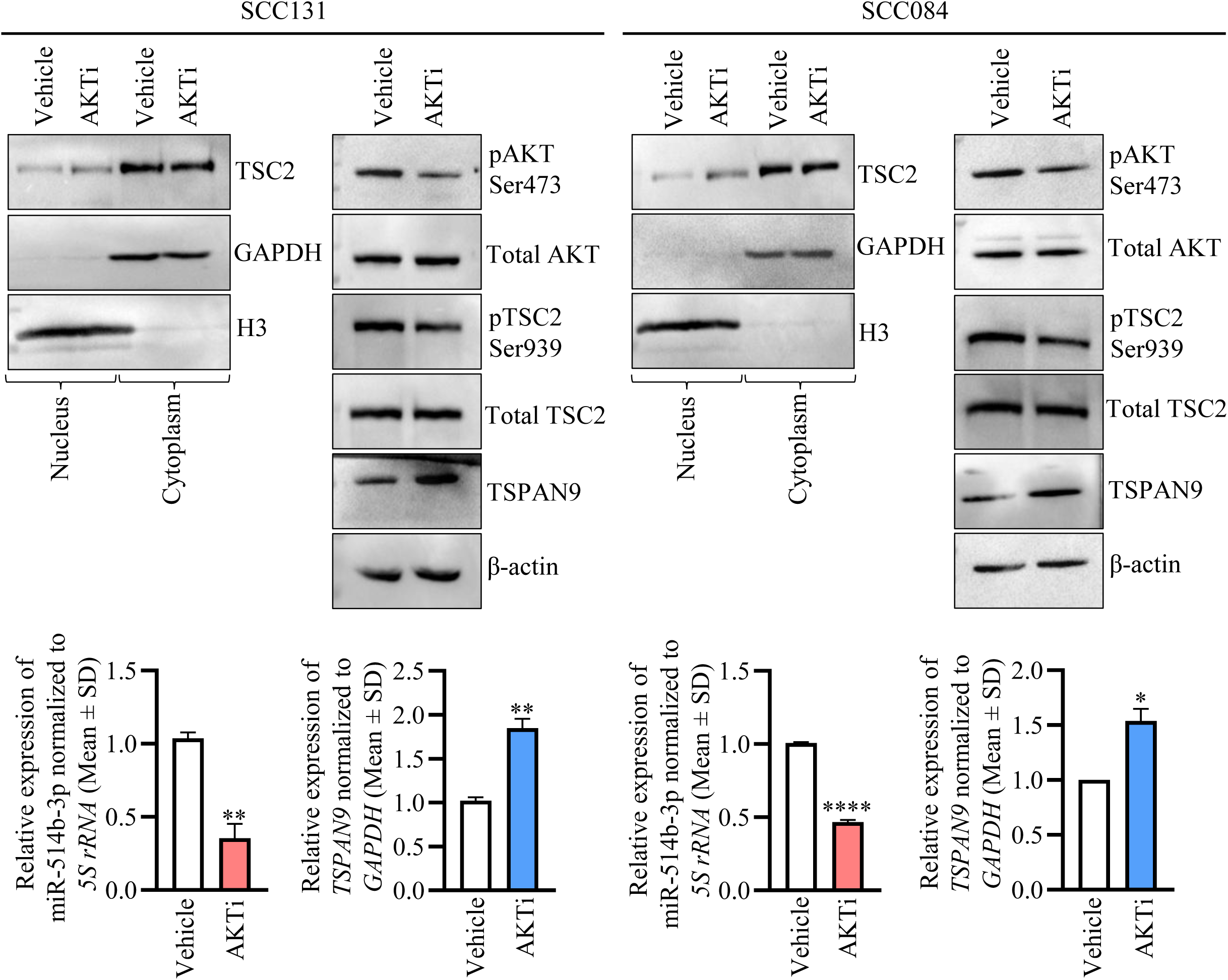
AKT regulates the miR-514b-3p/TSPAN9 axis via modulating TSC2 nuclear localization in SCC131 and SCC084 cells. Nuclear-cytoplasmic fractionations showing enhanced nuclear translocation of TSC2 following the AKTi treatment of OSCC cells. GAPDH and H3 were used as markers for cytoplasmic and nuclear fractions, respectively (upper panels). The levels of pAKT Ser473 and pTSC2 Ser939 decrease in AKTi treated cells as compared to those treated with the vehicle (upper panels). The miR-514b-3p expression decreases and *TSPAN9* expression increases after the AKTi treatment as compared to the vehicle-treated cells (upper and lower panels). β-actin was used as loading control. P-values were calculated with Unpaired student’s t-test with Welch correction. *Abbreviation:* AKTi, AKT inhibitor VIII.

The Western blot analysis of AKTi and DMSO-treated OSCC cells showed a sharp reduction in pAKT Ser473 and pTSC2 Ser939 levels in AKTi-treated cells compared to DMSO-treated cells, indicating an effective inhibition of AKT signalling by the AKT inhibitor VIII (Fig. 8; upper panels). The total AKT and total TSC2 levels remained unchanged in both treatments (Fig. 8; upper panels). Interestingly, we observed a significant reduction in the levels of miR-514b-3p in AKTi-treated cells as compared to those treated with DMSO, suggesting the nuclear TSC2-mediated inhibition of miR-514b-3p (Fig. 8; lower panels). Furthermore, we observed elevated levels of TSPAN9 in AKTi-treated cells compared with DMSO-treated cells, suggesting the miR-514b-3p-mediated regulation of *TSPAN9* (Fig. 8; upper and lower panels). These findings clearly demonstrated that AKT regulates the miR-514b-3p/TSPAN9 axis via modulating the nuclear localization of TSC2.

### Inhibition of MTORC1 does not affect the TSC2/miR-514b-3p/TSPAN9 axis

To rule out the possibility that TSC2 can regulate the miR-514b-3p/TSPAN9 axis indirectly via cytoplasm, we decided to inhibit MTORC1 (the primary downstream target of TSC2 in the cytoplasm) by treating OSCC cells with rapamycin and DMSO separately and examined its effect on the levels of miR-514b-3p and TSPAN9. As expected, we observed a sharp reduction in pS6K1 levels in rapamycin-treated cells as compared to those treated with DMSO, indicating an effective inhibition of MTORC1 (Fig. 9; upper panels). Further, we observed that the levels of miR-514b-3p and TSPAN9 remained unaltered in rapamycin and DMSO-treated cells, suggesting that the TSC2-mediated regulation of the miR-514b-3p/TSPAN9 axis is rapamycin-insensitive and hence independent of its MTORC1 function (Fig. 9).

**Fig. 9.**
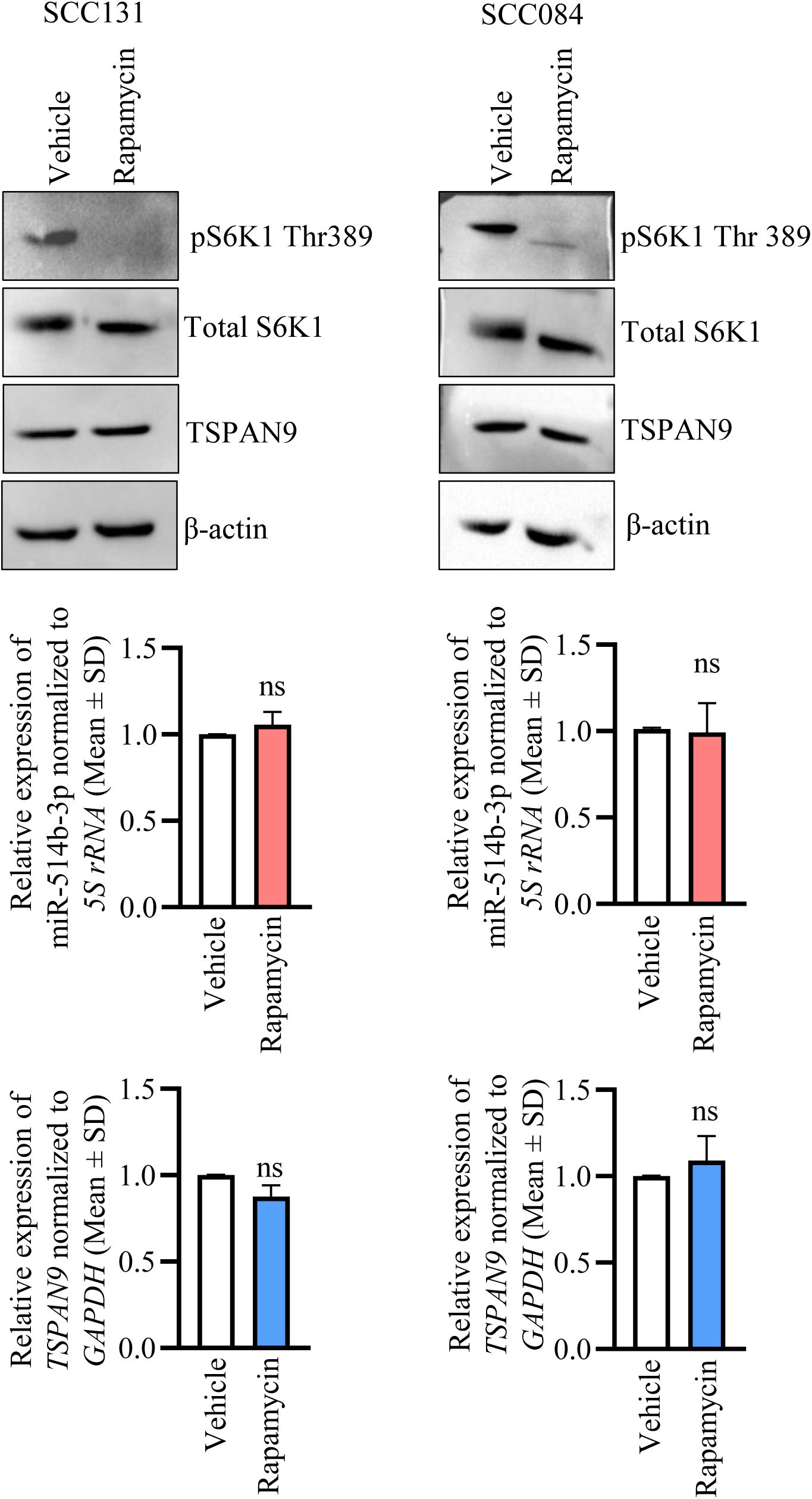
TSC2-mediated regulation of miR-514b-3p/TSPAN9 is independent of MTORC1 function in SCC131 and SCC084 cells. As expected, pS6K1 levels were sharply reduced in rapamycin-treated cells as compared to vehicle-treated cells (upper panels). Note, the levels of miR-514b-3p and TSPAN9 did not change in cells treated with rapamycin as compared to those treated with the vehicle control (upper and lower panels). β-actin was used as loading control. P-values were calculated with Unpaired student’s t-test with Welch correction.

## Discussion

The present study has demonstrated that TSC2 negatively regulates miR-514b-3p expression as a transcriptional repressor. In 1998, Henry et al. [15] discovered that TSC2 acts as a nuclear receptor co-regulator and mediates the transcriptional events led by the steroid receptor superfamily in a bidirectional manner. Specifically, TSC2 positively regulates the transcription mediated by VDR and PPRα, while repressing transcription mediated by the glucocorticoid receptor and ERα [15]. The direct interaction of TSC2 with VDR and ERα points towards its role as a nuclear receptor co-regulator [13,15].

The subsequent work from our laboratory identified *EREG* as the first transcriptional target of TSC2, demonstrating that TSC2 binds directly to the *EREG* promoter and suppresses its transcription [17]. While *EREG* represents a prototypical TSC2-regulated gene, it is unlikely to be the sole transcriptional target of TSC2. Given its nuclear localization and interaction with transcriptional machinery, it is reasonable to speculate that TSC2 regulates a broader repertoire of both coding and non-coding genes. In this context, Ogórek et al. [20] reported widespread alterations in miRNA expression following TSC2 loss in mouse embryonic fibroblasts, identifying 161 differentially expressed miRNAs, and demonstrating the role of TSC2 in miRNA biogenesis. However, this study attributed TSC2-mediated miRNA biogenesis to a cytoplasmic function of TSC2, mediated through the MTORC1 pathway [20]. In contrast, we sought to delineate the role of nuclear TSC2 in transcriptional regulation, with a particular focus on identifying miRNA genes that are regulated by TSC2 at the transcriptional level.

Our miRNA microarray analysis in SCC131 cells transfected with TSC2 siRNAs and Mock siRNAs led to the identification of 19 upregulated and 24 downregulated miRNAs, of which miR-514b-3p was one of the most robustly upregulated miRNAs (Fig. 1B, Table S1). The miR-514b-3p is encoded by the *MIR514B* gene located at chromosome Xq27.3 within an intergenic region, suggesting transcription from an independent promoter (UCSC Genome browser; https://genome.ucsc.edu/). Previous studies have reported context-dependent roles of miR-514b-3p in cancer, functioning either as an oncogene or a tumour suppressor [21–25]. Notably, miR-514b-3p has been shown to exert oncogenic functions in gastric cancer and tumour mycosis fungoides, where its expression is elevated [21,24]. Our observation of TSC2-mediated negative regulation of miR-514b-3p is consistent with these reports and suggests a putative oncogenic role for miR-514b-3p in OSCC cells (Fig. 1C and 1E).

Given that TSC2 represses *EREG* transcription via nucleus, we hypothesized that miR-514b-3p regulation might similarly depend on nuclear TSC2. York et al. [13] have demonstrated that TSC2 enters the nucleus via utilizing its NLS sequence near the C-terminus, and the deletion of this sequence restricts TSC2 nuclear entry. By selectively disrupting this NLS, we have showed that miR-514b-3p expression is significantly elevated in cells expressing the NLS-deleted TSC2 mutant compared with cells expressing the wild-type TSC2, suggesting that TSC2 negatively regulates miR-514b-3p levels in an NLS-dependent manner (Figs. 1D,1E, and S2). The deletion of NLS from the TSC2 suppressed the entry of exogenous TSC2 into the nucleus and resulted in hampering TSC2-mediated miR-514b-3p regulation.

To elucidate the underlying mechanism, we examined whether TSC2 directly regulates miR-514b-3p transcription by binding to the *MIR514B* promoter. Given the absence of neighbouring genes within a >10 Mbp region, it is likely that *MIR514B* expression is driven by its own independent promoter. Using a dual-luciferase reporter assay, we characterized the *MIR514B* promoter and demonstrated that TSC2 represses *MIR514B* promoter activity in a dose-dependent and NLS-dependent manner (Figs. 2A, 2B, and S4). This mode of regulation closely mirrors the previously described TSC2-mediated repression of the *EREG* promoter [17]. Further sequence analysis has revealed the presence of seven putative TSC2 binding sites (TBSs) within the *MIR514B* promoter (Fig. S5A and S5B). The subsequent ChIP-qPCR analysis has confirmed the direct binding of TSC2 to five out of the seven TBSs (TBS2, TBS3, TBS4, TBS5, and TBS6) in the *MIR514B* promoter, providing definitive evidence that TSC2 physically associates with the *MIR514B* promoter to repress its transcription (Fig. 2C). Notably, miR-514b-3p is the first identified non-coding (second overall after *EREG*) transcriptional target of TSC2. Hence, the nuclear TSC2 exerts its transcriptional function by binding to the promoter of its target genes, *EREG* and *MIR514B*, and negatively regulates their expression.

MiRNAs exert their downstream effects by interacting with the TS (target site) located within 3’UTRs, 5’UTRs, coding regions, and regulatory elements of their mRNA targets [26]. Given that *TSC2* is a classical tumour suppressor gene that negatively regulates miR-514b-3p expression, we sought to identify a downstream target of miR-514b-3p with a tumor-suppressive function. Through a combination of bioinformatics prediction and experimental validation, we identified *TSPAN9* (Tetraspanin 9) as a direct gene target of miR-514b-3p (Figs. 3 and S8; Table S4). TSPAN9 is a member of the tetraspanin family of proteins, a highly conserved group of transmembrane proteins that facilitate cellular signalling by assembling tetraspanin-enriched microdomains (TEMs) at the plasma membrane [27]. *TSPAN9* has been reported to act as a tumour suppressor in gastric and liver cancers, and as an oncogene in osteosarcoma, underscoring its context-dependent role in different cancers [28–31]. In OSCC cells, we observed a dose-dependent downregulation of TSPAN9 expression upon miR-514b-3p overexpression, indicating miR-514b-3p-mediated inhibition of *TSPAN9* (Fig. 3A and 3B). The bioinformatics analysis has revealed a single TS within the 3’UTR of *TSPAN9* (nucleotide positions 3432-3439, relative to TSS), complementary to the miR-514b-3p seed region (Fig. S8). The dual-luciferase reporter assay has confirmed that SR of miR-514b-3p interacts directly with the TS in the 3’UTR of *TSPAN9* in a sequence-specific manner (Fig. 3C). Consistently, the Western blot analysis has demonstrated that miR-514b-3p suppresses TSPAN9 protein expression through the direct binding to its 3’UTR in a sequence-specific manner (Fig. 3D). Importantly, the deletion of the target site from the 3’UTR of *TSPAN9* has abolished this interaction, resulting in loss of miR-514b-3p-mediated repression of TSPAN9 (Fig. 3C and 3D). Thus, the *TSPAN9* gene is another target of this miRNA, the others being *RBX* (Ring-box 1), *ARHGEF9* (Cdc42 guanine nucleotide exchange factor 9), *DUSP4* (Dual specificity phosphatase 4), *FZD4* (Frizzled class receptor 4), and *NTN1* (Netrin 1) [21–25].

Functionally, our results have demonstrated that miR-514b-3p promotes and *TSPAN9* suppresses proliferation of OSCC cells, consistent with their respective oncogenic and tumour suppressive roles (Figs. 5 and S10). These findings align with earlier observations in gastric cancer, where miR-514b-3p functions as an oncomiR and *TSPAN9* acts as a tumour suppressor [24,28,29,32]. Furthermore, we have shown that miR-514b-3p enhances cell proliferation and anchorage-independent growth, while inhibits apoptosis of OSCC cells, in part, by targeting the 3’UTR of *TSPAN9* in a sequence-specific manner (Figs. 6 and S11). It is noteworthy that by suppressing *TSPAN9* expression and hence its tumour suppressor function, miR-514b-3p exhibits its oncogenic properties via regulating these cancer hallmarks. Disruption of the miR-514b-3p/TSPAN9 interaction has abrogated these effects, reinforcing the functional relevance of this regulatory interaction in governing key cancer hallmarks (Figs. 6 and S11).

Consistent with its role as a transcriptional repressor of miR-514b-3p, TSC2 positively regulates TSPAN9 expression in an NLS-dependent manner (Fig. 4). Given the direct binding between TSC2 and the *MIR514B* promoter, as well as miR-514b-3p and *TSPAN9*, we reasoned that TSC2 should regulate *TSPAN9* via miR-514b-3p regulation. Thus, tumour suppressor *TSC2* inhibits the oncogenic miR-514b-3p, while miR-514b-3p suppresses the expression of another tumour suppressor *TSPAN9*, establishing a regulatory axis TSC2/miR-514b-3p/TSPAN9. Collectively, our study identified a novel, nucleus-mediated TSC2/miR-514b-3p/TSPAN9 axis in OSCC cells.

We next investigated whether this newly found axis intersects with the canonical PI3K-AKT-MTOR pathway in the cytoplasm. TSPAN9 has previously been shown to suppress the PI3K-AKT-MTOR signalling by promoting degradation of the PI3K p55 subunit in gastric cancer [29]. In agreement with this observation, we observed upregulation of p55, pAKT, and pS6K1 levels following miR-514b-3p overexpression and downregulation of p55, pAKT, and pS6K1 levels following TSPAN9 overexpression in OSCC cells (Fig. 7). The miR-514b-3p overexpression caused reduced levels of TSPAN9, leading to the upregulation of p55 and subsequent activation of AKT and S6K1 (Fig. 7). Notably, the alteration in TSC2 levels regulated the miR-514b-3p/TSPAN9 axis, whereas alteration in miR-514b-3p/TSPAN9 levels further modulated the PI3K-AKT-MTOR pathway, suggesting the existence of a positive feedback loop (Fig. 7). In addition, the miR-514b-3p-mediated activation and TSPAN9-mediated inhibition of the PI3K-AKT-MTOR pathway further ensured their respective oncogenic and tumour suppressor potential in OSCC cells. This is particularly noteworthy that TSC2 negatively regulates the PI3K-AKT-MTOR pathway via RHEB inhibition in the cytoplasm and as a transcriptional repressor of oncogenes, *EREG* and *MIR514B* in the nucleus.

AKT is known to regulate the nuclear-cytoplasmic shuttling of TSC2 through phosphorylation at Ser939 and Thr1462, thereby restricting its nuclear functions [19]. In line with this, pharmacological inhibition of AKT with AKTi (AKT inhibitor VIII) in OSCC cells resulted in downregulation of miR-514b-3p and upregulation of TSPAN9 levels (Fig. 8). These observations can be attributed to the enhanced translocation of TSC2 in the nuclear compartment following AKT inhibition, which leads to transcriptional suppression of miR-514b-3p, resulting in the elevated TSPAN9 levels (Fig. 8). These results further underscored the inter-dependence between the canonical PI3K-AKT-MTOR signalling and the newly identified TSC2/miR-514b-3p/TSPAN9 axis. To enunciate that the regulation of miR-514b-3p/TSPAN9 axis is independent from cytoplasmic role of TSC2, we treated OSCC cells with rapamycin, a widely known inhibitor of the MTORC1 activity [11]. Notably, inhibition of MTORC1 activity did not alter miR-514b-3p and TSPAN9 levels, confirming that the regulation of the miR-514b-3p/TSPAN9 axis is MTORC1-independent and is not a cytoplasmic function of TSC2 (Fig. 9).

Taken together, this study identifies miR-514b-3p as the first non-coding transcriptional target of TSC2 and delineated the molecular mechanism underlying its regulation. The study establishes *TSPAN9* as a downstream target of miR-514b-3p and discovered a novel nucleus-driven TSC2-miR-514b-3p-TSPAN9 axis in OSCC cells. Further research on the oncogenic properties of miR-514b-3p and the tumour suppressor properties of TSPAN9 might benefit therapeutic interventions in OSCC and other malignancies.

## Materials and Methods

### Plasmid constructs

The details of plasmid constructs generated for the present study are summarized in Table S5. The pPROM-MIR514B, pMIR514B, p3’UTR-TSPAN9, p3’UTR-RBX, and pTSPAN9 constructs were generated using human genomic DNA or cDNA as a template and gene specific primers (Table S5) by standard laboratory protocols [33]. The p3’UTR-TSPAN9-M was generated by site-directed mutagenesis using specific primers (Table S5). pTSPAN9-3’UTR and pTSPAN9-3’UTR-M were generated by cloning *TSPAN9* 3’UTR and *TSPAN9* 3’UTR-M respectively downstream to pTSPAN9. All the constructs were subjected to Sanger sequencing on an ABI PRISM A310-automated sequencer (Thermo Fisher Scientific, Waltham, MA, USA) to confirm directionality of the inserts and error-free sequences.

### Cell culture and transient transfections

Human oral squamous cell carcinoma cell lines, UPCI-SCC-131 and UPCI-SCC-084, were a kind gift from Prof. Susanne M. Gollin, University of Pittsburgh, PA, USA. These cell lines were cultured in DMEM (Dulbecco’s Modified Eagle Medium, high glucose), supplemented with 10% FBS (Fetal Bovine Serum) and 1X Antibiotic-Antimycotic solution (all from Sigma-Aldrich, St. Louis, MO, USA) at 37⁰C in 5% CO_2_.

SCC131 or SCC084 cells were seeded at a density of 2 × 10^6^ cells per well in a 6-well plate and transiently transfected with appropriate plasmid construct(s) using the Lipofectamine 2000 transfection reagent (Thermo Fisher Scientific, Waltham, MA, USA) as per the manufacturer’s instructions. After 48 hr, cells were harvested for RNA and protein isolation as per experimental plan.

### siRNA mediated TSC2 knockdown

A SMART pool of four siRNAs targeting *TSC2* transcript (Cat# L-003029-00-0005) and a non-targeting pool of four siRNAs (Mock) (Cat# D-001810-10-05) were commercially purchased from the GE Healthcare Dharmacon, Lafayette, CO, USA (Table S6). SCC131 cells were seeded at a density of 2 × 10^6^ cells per well in a 6-well plate and transiently transfected either with 100 nM Mock siRNAs or TSC2 siRNAs pool separately, using Lipofectamine 2000 following the manufacturer’s instructions. After 48 hr, cells were harvested for small RNA, total RNA, and total protein isolation as per experimental plan.

### Microarray analysis

The small RNA isolated from Mock and TSC2 siRNAs-transfected cells (3 replicates each) were subjected to the microRNA microarray analysis (Genotypic technology, Bengaluru, India). The microRNA microarray was performed on SurePrint G3 8 × 60k Human miRNA microarray platform having 2,549 mature miRNAs probes (AMADID 70156; Agilent Technologies, Santa Clara, CA, USA). The miRNA microarray data was analyzed to identify differentially regulated miRNAs in TSC2 depleted cells. A list of differentially regulated miRNAs following TSC2 knockdown is given in Table S1.

### Bioinformatics analysis to identify miR-514b-3p targets

We have performed bioinformatics analysis to predict the gene targets for miR-514b-3p. The bioinformatics tools used for the gene target prediction of miR-514b-3p were Agilent GeneSpring (https://shorturl.at/4dJHb), miRDB [34], TargetScanHuman8.0 [35], microT-CDS [36], and STarMirDB [37].

### RNA isolation and cDNA synthesis

The total RNA was isolated using TRI-Reagent as per the manufacturer’s protocol (Sigma-Aldrich, St. Louis, MO, USA) and quantified using a NanoDrop 1000 spectrophotometer (Thermo Fisher Scientific, Waltham, MA, USA). The first-strand cDNA synthesis was performed using a Verso cDNA synthesis kit (Thermo Fisher Scientific, Waltham, MA, USA) and 1-2 µg of total RNA as a template.

### Reverse transcription-quantitative PCR

The transcript level of miR-514b-3p was determined by the RT-qPCR-based miR-Q assay [38], whereas for all other transcripts, the expression levels were determined by conventional RT-qPCR. The details of the RT-qPCR primers are given in Table S2. The RT-qPCR was performed using a DyNAmo ColorFlash SYBR Green qPCR kit (Thermo Fisher Scientific, Waltham, MA, USA) in a QuantStudio 3 PCR system (Thermo Fisher Scientific, Waltham, MA, USA) as per the manufacturer’s instructions. The RT-qPCR data was analyzed to calculate the relative gene expression using the following formula: ΔC_t (gene)_ = C_t (gene)_ – C_t (normalizing control)_, followed by calculating the fold change using 2^(−ΔΔC_t_), where C_t_ represents the cycle threshold value. 5S *rRNA* and *GAPDH* were used as a normalization controls. Each bar represents the relative fold change expression of a gene of interest normalized to its housekeeping gene and is an average of 3 biological replicates.

### Western blot analysis

The total protein was isolated using the CelLytic M Cell Lysis reagent (Sigma-Aldrich, St. Louis, MO, USA), resolved on SDS-PAGE, and transferred to a PVDF membrane (Cytiva, Marlborough, MA, USA). The membrane was probed with an appropriate primary antibody and visualized by a ChemiDoc imaging system equipped with a CCD camera (Bio-rad, Inc, Hercules, CA, USA) using an Immobilon Western chemiluminescent HRP substrate (Merck-Millipore, Darmstadt, Germany). Details of antibodies and their dilutions used for the Western blot analysis are mentioned in Table S7.

### Dual-luciferase reporter assay

For the dual-luciferase reporter assay, 50,000 SCC131 or SCC084 cells were seeded in a 24-well plate and were transiently transfected with appropriate plasmid construct(s) as per the experiment requirement. The assay was performed 48 hr post transfection using a Dual-Luciferase^®^ Reporter (DLR™) Assay System kit (Promega, Madison, WI, USA) as per the manufacturer’s protocol. The pRL-TK control vector was co-transfected to normalize the transfection efficiency. Each bar is an average of 3 biological replicates and represents relative light units (RLU).

### ChIP-qPCR

The ChIP-qPCR was performed using a SimpleChIP^®^ Enzymatic Chromatin IP kit (Magnetic Beads) (Cell Signaling Technology, Danvers, MA, USA) as per manufacturer’s instructions. Briefly, 2 × 10^6^ SCC131 cells per 60 mm dish were seeded. Upon reaching ∼90% confluency, the cells were cross-linked using 37% formaldehyde per dish (Sigma-Aldrich, St. Louis, MO, USA), following which the chromatin was isolated for three IP (immunoprecipitation) reactions using TSC2, IgG, and H3 antibodies. For each ChIP reaction, 5 to 10 µg of digested, cross-linked chromatin was used. After pulldown, all IP reactions were reverse cross-linked, and DNA was purified from each sample, which was used as a template for the ChIP-qPCR analysis. The primers were designed to amplify the regions spanning TBSs (TSC2 binding sites) to check the enrichment of TSC2 on the *MIR514B* promoter (Table S3). The amplification in the IgG-ChIP DNA sample was used as a negative control and was compared to analyse the fold enrichment in TSC2-ChIP DNA. For the positive control, the *RPL30* gene was amplified with H3-ChIP DNA as a template.

### Cell proliferation assays

The effect of miR-514b-3p, TSC2 and TSPAN9 on proliferation of SCC131 and SCC084 cells was studied by the trypan blue dye exclusion [39] and the BrdU cell proliferation assays [40]. For, trypan blue dye exclusion assay, 2 × 10^6^ cells per well in a 6-well plate were seeded in triplicates and transiently transfected with appropriate plasmid construct(s), using the Lipofectamine 2000 transfection reagent. On the day of counting, cell suspension was mixed with 0.4% trypan blue dye (Sigma-Aldrich, St. Louis, MO, USA), loaded onto a Neubauer haemocytometer (HyClone, Logan, UT, USA), and counted using an Olympus CKX4 inverted microscope (Olympus Optical Co., Tokyo, Japan). The total cell number in the original suspension was calculated by the following formula: number of cells/mL = average of cells in four quadrants × dilution factor (10) × 10^4^. The live cells were counted every 24 hr for four days, and viable cell number was plotted as a function of time.

For the BrdU cell proliferation assay, 2,000 cells per well were seeded in quadruplicate in a 96-well plate (3 days, 3 total plates). The cell proliferation was assessed on Day 2, Day 4, and Day 6, using a BrdU cell proliferation assay kit as per the manufacturer’s instructions (Merck-Millipore, Darmstadt, Germany).

### Caspase-3 activation assay

The apoptosis was measured using a Caspase-3 assay kit (Abcam, Cambridge, UK) as per the manufacturer’s instructions. Briefly, 2 × 10^6^ cells per well were seeded in a 6-well plate and transfected using Lipofectamine 2000 with appropriate plasmid construct(s). Post 48 hr, 1 µL of FITC-DEVD label was added to each cell suspension, following which its fluorescence intensity was measured (Ex: 485 nm; Em: 535 nm) using a plate reader (Tecan Group Ltd, Mannedorf, Switzerland).

### Anchorage independence assay

The ability of cells to grow in an anchorage independent manner was assessed by the soft agar colony formation assay [41]. After transfecting cells with appropriate constructs, they were harvested and 10,000 cells were plated in 1 mL of 0.5% Noble Agar (BD Difco, Franklin Lakes, NJ, USA) diluted with culture media in a 35 mm dish. After three weeks, colonies were imaged with a 2.5X objective lens using Leica Inverted Microscope Dmi1 (Leica Microsystems, Wetzlar, Germany) and counted using an image analysis software (Fiji; https://imagej.net/ij/).

### AKT inhibitor VIII and rapamycin treatments

For both treatments, 2 × 10^6^ SCC131 or SCC084 cells were seeded in a 6-well plate and treated with 58 nM AKT inhibitor VIII (AKTi) for 24 hr or 100 nM rapamycin (MedChemExpress, Princeton, NJ, USA) for 6 hr respectively [42]. DMSO was used as vehicle control in both treatments. After appropriate incubation time, the cells were harvested for total RNA and protein isolation for subsequent experiments.

### Statistical analysis

All the experimental data (bar graphs and line diagrams) were plotted using the GraphPad Prism 8 software (https://www.graphpad.com/). The comparison between any two experimental data sets was performed by calculating the statistical significance using Unpaired student’s t-test with Welch correction. The comparison among multiple data sets was carried out by performing one-way ANOVA, followed by Tukey’s multiple comparison test. The difference among data sets was considered significant when p-values were ≤0.05 (*), <0.01 (**), <0.001 (***), <0.0001 (****) or non-significant (ns) when the p-value was >0.05.

## Author contributions

SG planned and performed all the experiments, analyzed data, wrote the manuscript and made all figures. NM prepared TSC2 knocked down samples for miRNA microarray. MK assisted in ChIP and cancer hallmark assays. AK supervised the project, procured funding and resources and edited the final version of the manuscript. All authors reviewed and approved the final version of the manuscript.

## Data availability statement

All data are available in the manuscript or available upon request.

