## Supplementary figures and Tables for "Nuclear TSC2 drives miR-514b-3p transcription to regulate PI3K-AKT-MTOR signalling: Implications for OSCC pathogenesis"

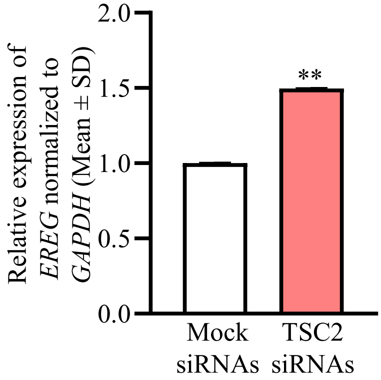


**Fig. S1. Validation of TSC2 knockdown in SCC131 cells.** Upregulation of the *EREG* transcript level in TSC2 knockdown cells as compared to those transfected with the Mock siRNAs. P-value was calculated with Unpaired student's t-test with Welch correction.

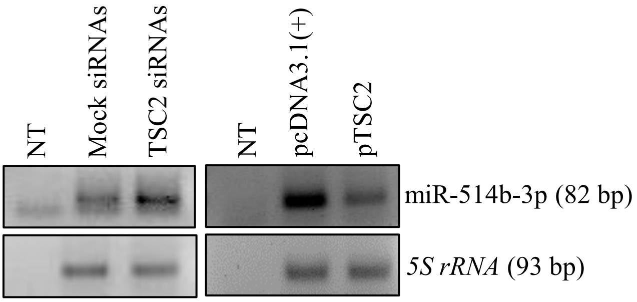


**Fig. S2. TSC2 negatively regulates the miR-514b-3p expression.** The upregulation of miR-514b-3p in TSC2 siRNAs-transfected SCC131 cells as compared to those transfected with Mock siRNAs by RT-PCR (left panels). The downregulation of miR-514b-3p in pTSC2-transfected cells as compared to those transfected with pcDNA3.1(+) by RT-PCR (right panels). *Abbreviation*: NT, No template control.

(A)

**-2575**
**5'**CCCTCTCATGGGCATTTTCTGTCATATATTTTTGTTATCTGTAAAGACTACACTATCTTTTTTTTGTACATATTATGATTTTCTGTAAAACAGGA
TAGTTAAGATCAAATATTGTATTCTGGATAATAACCACTTCCTCTGAGGAATTATATTTTTTTTGTTTCTTTGTTTTGTTTTTGAGATGGAGTCTC
GCTCTATCACCCAGGTTGGAGTGCAGTGGCGTGATCGCGGCTCACTGCGACCTCTGCCTCCTGGGTTCAAGCGATTCTCCTGCCTCAGCCTCCT
GAGTACCTGGGACTACAGGCGCCCGCCACCGCGCCCAGCTAATTTTTGTATTTTTAGTAGAGACGGGGTTTCACCATGTTGGCCAGGATGGTCT
CAATCTCCTGACCTGGTGATCTACCCACCTCAGCCTCCCAAAGTGCTGGGCTTTACAGACGTGAGCCACCGCACCTGGCTTTTACTGTTGTTGTT
GTTGTCTGTTTTTAATTTGTTTATTGAATTGGCTGGAGTTTATTTACTTATCTGCTGTGAATTTTCTTATACTGTTCTTTAGAGAGCACAGCCTTT
GTATGCACATAGTAACTCTAAGATGACAGTGATTTTAGTAGCATTCTTTTTGTTTCTTTTCCTGATCTCTCTGTCCAGCTAGTGGTCTCTGTGTAT
CACACAGAGATGTTAGGTTTTCTCACTGAATGCTGACTTTCTCTATCGTTTTTTTTGAATTTTAAGGGAAAAACGTATCAGAGTTTGATCAAAAT
AATTTAAGCCACTTGGAACACGTTGTTTTAAAGGCAAGTCTTTCAGGTTTGTTCTGATACAGGAAGGCTCTTATTAGCTGTCAGTTTATCGGAT
ACTCTGGTAAATAAGCTGGCCTGTGGTTTAGCTTGTTGCTCTCATACAGCTACCAAATCCATCTTACTTACTTACCTCCAAAATCTGTGTTGTTA
CCTGGAGCATCGTTAAGCCTGAGCTTTCCCTGACTCTGCTATAAATAAAGTCCTATTTTTTTTTTTAGGAGAGCTATAATACCATCTTTTATTCA
AACTCACTTCTCACACTGGGAAAAAGAATCTCTGACTCACTGCTCTCAAGCTTTGGGAAGGTGCAGCGACCTGCTTCTTTCACAACACTCCTGC
TTTATGTACAGAGTGAAGATTGGTAGTCGTAGATTTTGGTCTTCTCAGCCTGCTCATTCCAGGGTTGAAACTCTGTTTTATGATTATTCTGGGGT
GATGATGATCAGAGACCCCATATTCTCAACCTGCCAAAACCACACTAAAGCCTACACCCTACGAAAGTGAGCTGAGTGTACGAACAGTGCCC
CTAACATCTGTACCTGTTGCTTAGCCACAGCAACACAAAATTAAGGGTAGGGTGCATGAGAAAGGCTGGGAACTTGTTTCTTCCATTAAGATA
CCATAATTTTTGACTGGTAGCAACTTCACATTTTAGGCTACTTACAATCACCTCTTTCTTCCATCCCAGTTCTCTGACATCCATTTACCATATTTG
TGTTACTATATTTTGGTGTCCCAGATTTAGTTCATCCAGGAAAATGAAGACACATAGTTGCATGTTTCCAAATGGAGAAAAATGCAAACAAAA
ATGCTTGAAAAGACCACGCTCGAGGGAAAGACTCAGAAATCTCACAAACTGGAAGAAGTTTAATGGCATCCATAGATGCCAAGAACATTTTC
TGTTTTGCAGTTCATTCCTGGTAAGTAAAATATTTTTATTTTTGCTCTAGGATTTAAGGAGAAACAGACATTTCTACTCTCCATCACATCAGTGA
TTACCATCCTATCCTAGAATCTTGTGTGTTTTCCCTGCTTATTCTGTGGTTCAGGAGTTGAGGGATGTAACTCAGGTAGAGGGACTCATCTTCAG
GTGACGAGCATAGATGACATAGAGTATATTTTCCTGTTAACTATTAGACCTGAGGAATGTAAACTGCTTTCTAACACTTTATAATAATGGAAA
AAATAAACAGGATAAGACATGCAAGGATTTTTCTTCCGTGCACAAGAGAGTCTAGCAGAAAAGCTTAATGGTTCATAGTTCTGAAAATTGCCA
TGTCATAACTCCAAACCATGATTGGACAACACAGGAAAGGATGGATGAAGAAAGACAGTGTAGTGCATGCTACAGAGGCCCTTTGTATGAAC
ACCAGTCACCAATCAGATTGAAAACAATGATGTGTTTTTGAAAGATGCATGTTGGTCAAAGAGTACAAACTTTGAGTTACGTAATAGATAAGC
TCTGGCTATCTAATGTATAGCATAGTGATAATAATGCTGTAATTGTTTACTGGAAATTTGATAAGAGAGCAGATTTTAAGTGTCCTCACCACAC
CCCCACAACACACACACAAATGGTAACTAAGAGCGGTGAATGATGTGTTAGTTAATTTTACTATGGTAATCAGTACGCGATGTATATACATAT
CAAATCACATTGCACACCTTGAATATATACAAATTTTGTTTGTCAATTACATAT
**General TF binding site** **TSS +1**
TTTTAAAATATAAAAAGAAAGGTGCATGAGTCAGTCATTAGGCTGGGCTGCCATCCTTAAGTGTTCTATTCTTCATGTGGTACTCTTCTCAAGA
GGGAGGCAATCATGTGTAATTAGATATGATTGACACCTCTGTGAGTGGAGTAACACATGATATATACATCATATGACATGGCATGGTGGATAC
ACGGGGCGTGTTAAGTTGCAAGCAGTATAATACTTTCCCTAGAATGCAGACATTTCTACTGACACCCTCCCTGTTCTGGACTAGCTCAGTTAAT
TCTCTAAATTAATTATCAGACGTAAGACCTAGAAACTGAAAG **3'**
**+250**

(B)

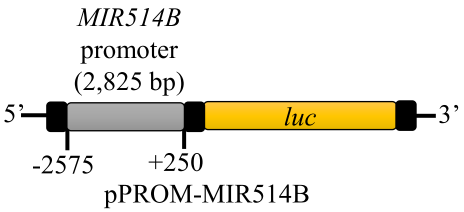


**Fig. S3. The predicted promoter sequence for the *MIR514B* gene.** (A) The 2,825 bp long putative promoter sequence of *MIR514B* as predicted by the DBTSS and the Promoter 2.0 databases. The general TF binding site is highlighted in yellow colour, whereas the TSS is highlighted in red colour. (B) A diagrammatic representation of the *MIR514B* promoter construct pPROM-MIR514B. *Abbreviation*: TF, transcription factor; and TSS, transcription start site.

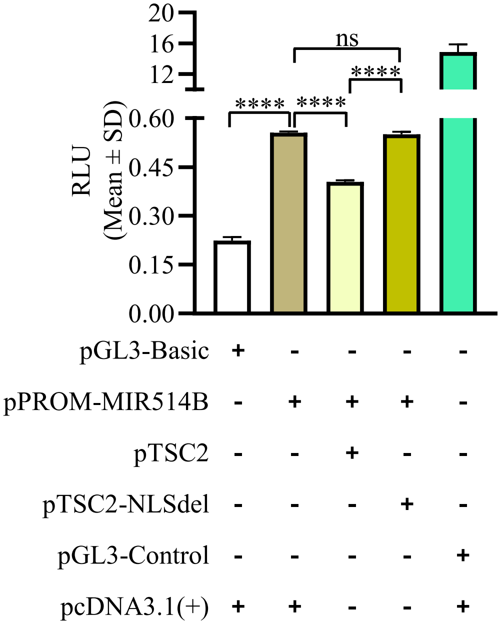


**Fig. S4. TSC2 negatively regulates the *MIR514B* promoter activity.** The SCC084 cells co-transfected with pPROM-MIR514B and pTSC2 showed a significantly reduced luciferase activity as compared to those co-transfected with pPROM-MIR514B and the vector pcDNA3.1(+), whereas the cells with pPROM-MIR514B and pTSC2-NLSdel showed a significantly increased luciferase activity as compared to those co-transfected with pPROM-MIR514B and pTSC2 (N=3). P-values were calculated with one way ANOVA with Tukey's multiple comparison test. *Abbreviation*: RLU, Relative light unit.

(A)

**-2575**
**5’**CCCTCTCATGGGCATTTTCTGTCATATATTTTTGTTATCTGTAAAGACTACACTATCTTTTTTTTGTACATATTATGATTTTCTGTAAAACAGGATAGTTAAG
ATCAAATATTGTATTCTGGATAATAACCACTTCCTCTGAGGAATTATATTTTTTTTGTTTCTTTGTTTTGTTTTTGAGATGGAGTCTCGCTCTATCACCCAGGT
TGGAGTGCAGTGGCGTGATCGCGGCTCACTGCGACCTCTGCCTCCTGGGTTCAAGCGATTCTCCTGCCTCAGCCTCCTGAGTACCTGGGACTACAGGCGCCC
GCCACCGCGCCCAGCTAATTTTTGTATTTTTAGTAGAGACGGGGTTTCACCATGTTGGCCAGGATGGTCTCAATCTCCTGACCTGGTGATCTACCCACCTCA
GCCTCCCAAAGTGCTGGGCTTTACAGACGTGAGCCACCGCACCTGGCTTTTACTGTTGTTGTTGTTGTCTGTTTTTAATTTGTTTATTGAATTGGCTGGAGTTT
ATTTACTTATCTGCTGTGAATTTTCTTATACTGTTCTTTAGAGAGCACA**GCCTTTG**TATGCACATAGTAACTCTAAGATGACAGTGATTTTAGTAGCATTCTT
TTTGTTTCTTTTCCTGATCTCTCTGTCCAGCTAGTGGTCTCTGTGTATCACACAGAGATGTTAGGTTTTCTCACTGAATGCTGACTTTCTCTATCGTTTTTTTTG
AATTTTAAGGGAAAAACGTATCAGAGTTTGATCAAAATAATTTAAGCCACTTGGAACACGTTGTTTTAAAGGCAAGTCTTTCAGGTTTGTTCTGATACAGG
AAGGCTCTTATTAGCTGTCAGTTTATCGGATACTCTGGTAAATAAGCTG**GCCTG**TGGTTTAGCTTGTTGCTCTCATACAGCTACCAAATCCATCTTACTTACT
TACCTCCAAAATCTGTGTTGTTACCTGGAGCATCGTTAA**GCCTG**AGCTTTCCCTGACTCTGCTATAAATAAAGTCCTATTTTTTTTTTTAGGAGAGCTATAAT
ACCATCTTTTATTCAAACTCACTTCTCACACTGGGAAAAAGAATCTCTGACTCACTGCTCTCAAGCTTTGGGAAGGTGCAGCGACCTGCTTCTTTCACAACA
CTCCTGCTTTATGTACAGAGTGAAGATTGGTAGTCGTAGATTTTGGTCTTCTCA**GCCTG**CTCATTCCAGGGTTGAAACTCTGTTTTATGATTATTCTGGGGTG
ATGATGATCAGAGACCCCATATTCTCAACCTGCCAAAACCACACTAAAGCCTACACCCTACGAAAGTGAGCTGAGTGTACGAACAGTGCCCCTAACATCTG
TACCTGTTGCTTAGCCACAGCAACACAAAATTAAGGGTAGGGTGCATGAGAAAGGCTGGGAACTTGTTTCTTCCATTAAGATACCATAATTTTTGACTGGT
AGCAACTTCACATTTTAGGCTACTTACAATCACCTCTTTCTTCCATCCCAGTTCTCTGACATCCATTTACCATATTTGTGTTACTATATTTTGGTGTCCCAGAT
TTAGTTCATCCAGGAAAATGAAGACACATAGTTGCATGTTTCCAAATGGAGAAAAATGCAAACAAAAATGCTTGAAAAGACCACGCTCGAGGGAAAGACT
CAGAAATCTCACAAACTGGAAGAAGTTTAATGGCATCCATAGATGCCAAGAACATTTTCTGTTTTGCAGTTCATTCCTGGTAAGTAAAATATTTTTATTTTT
GCTCTAGGATTTAAGGAGAAACAGACATTTCTACTCTCCATCACATCAGTGATTACCATCCTATCCTAGAATCTTGTGTGTTTTCCCTGCTTATTCTGTGGTT
CAGGAGTTGAGGGATGTAACTCAGGTAGAGGGACTCATCTTCAGGTGACGAGCATAGATGACATAGAGTATATTTTCCTGTTAACTATTAGACCTGAGGAA
TGTAAACTGCTTTCTAACACTTTATAATAATGGAAAAAATAAACAGGATAAGACATGCAAGGATTTTTCTTCCGTGCACAAGAGAGTCTAGCAGAAAAGCT
TAATGGTTCATAGTTCTGAAAATT
**GCCATG**TCATAACTCCAAACCATGATTGGACAACACAGGAAAGGATGGATGAAGAAAGACAGTGTAGTGCATGCTACAGAG**GCCCTTTG**TATGAACACC
AGTCACCAATCAGATTGAAAACAATGATGTGTTTTTGAAAGATGCATGTTGGTCAAAGAGTACAAACTTTGAGTTACGTAATAGATAAGCTCTGGCTATCT
AATGTATAGCATAGTGATAATAATGCTGTAATTGTTTACTGGAAATTTGATAAGAGAGCAGATTTTAAGTGTCCTCACCACACCCCCACAACACACACACA
AATGGTAACTAAGAGCGGTGAATGATGTGTTAGTTAATTTTACTATGGTAATCAGTACGCGATGTATATACATATCA
**General TF binding site**
TCACATTGCACA**CCTTG**AATATATACAAATTTTGTTTGTCAATTACATATTTTTAAAATATAAAAAGAAAGGTGCA
**TSS +1**
TGAGTCAGTCATTAGGCTGGGCTGCCATCCTTAAGTGTTCTATTCTTCATGTGGTACTCTTCTCAAGAGGGAGGCAATCATGTGTAATTAGATATGATTGAC
ACCTCTGTGAGTGGAGTAACACATGATATATACATCATATGACATGGCATGGTGGATACACGGGGCGTGTTAAGTTGCAAGCAGTATAATACTTTCCCTAG
AATGCAGACATTTCTACTGACACCCTCCCTGTTCTGGACTAGCTCAGTTAATTCTCTAAATTAATTATCAGACGTAAGACCTAGAAACTGAAAG **3’ +250**

(B)

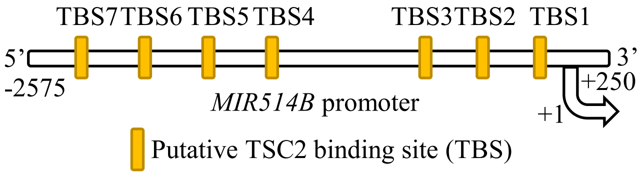


**Fig. S5. Putative TSC2 binding sites (TBSs) in the *MIR514B* promoter.** (A) Seven TBSs predicted in the 2,825 bp long *MIR514B* promoter sequence are highlighted in blue colour. The general TF binding site is highlighted in yellow colour, and the TSS is highlighted in red. (B) A schematic representation of the *MIR514B* promoter with seven predicted TBSs. TSS is numbered from nucleotide position +1. *Abbreviations*: TF, Transcription factor; TSS, Transcription start site; and TBS, TSC2 binding site.

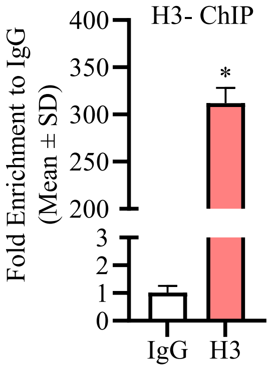


**Fig. S6. H3 ChIP as an internal positive control.** The H3 protein showed binding to the *RPL30* region and was served as an internal control. P-value was calculated with Unpaired student's t-test with Welch correction.

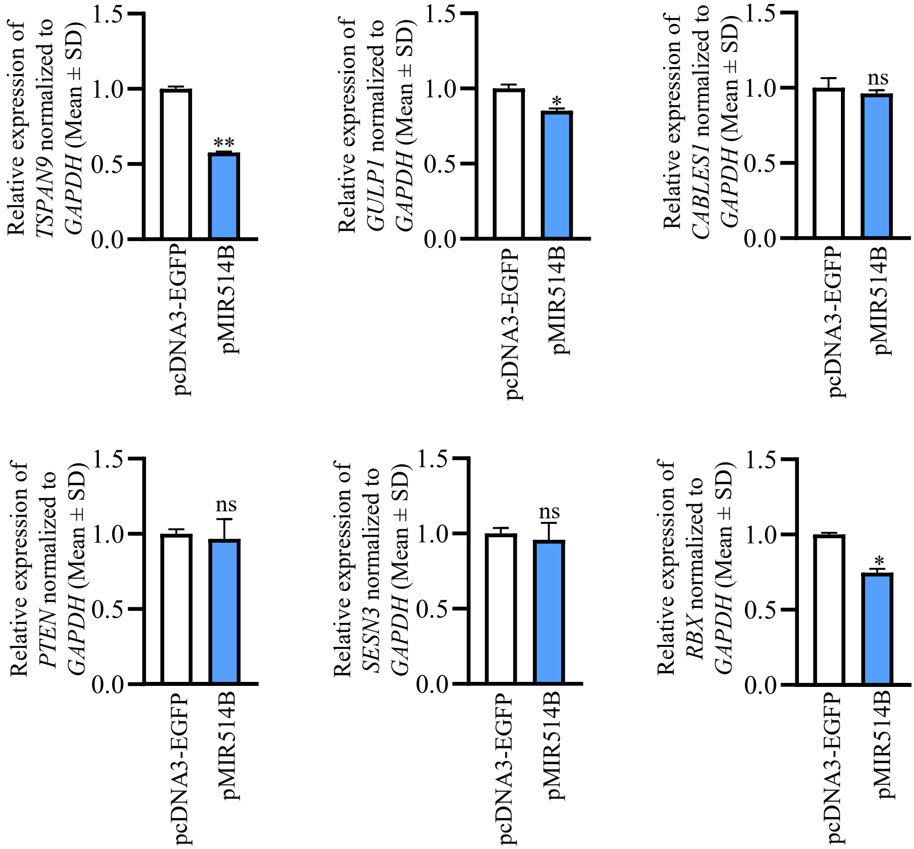


**Fig. S7. The effect of miR-514b-3p overexpression on the transcript levels of potential target genes of miR-514b-3p in SCC131 cells.** Note, overexpression of miR-514b-3p causes a significant downregulation of transcript levels of *TSPAN9* and *GULP1* in cells transfected with pMIR514B compared to those transfected with pcDNA3-EGFP, whereas the transcript levels of *CABLES*, *SESN3*, and *PTEN* remain unchanged in cells transfected with pMIR514B compared to those transfected with pcDNA3-EGFP. As expected, the transcript level of a known target gene *RBX* of miR-514b-3p was found to be significantly downregulated in cells transfected with pMIR514B compared to those transfected with pcDNA3-EGFP. P-values were calculated with Unpaired student's t-test with Welch correction.

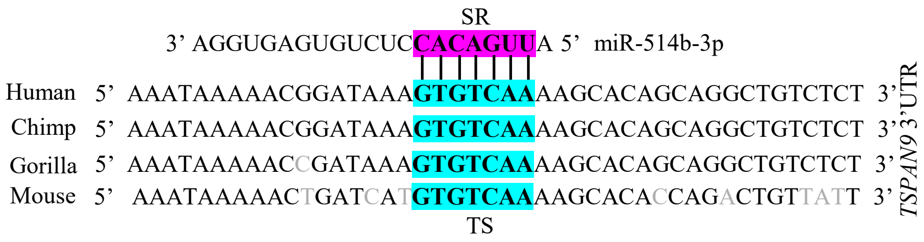


**Fig. S8. Conservation of TS in the 3’UTR of *TSPAN9* across species.** Note, the base complementarity of SR of miR-514b-3p with TS of *TSPAN9*. *Abbreviations*: TS, target site; and SR, seed region.

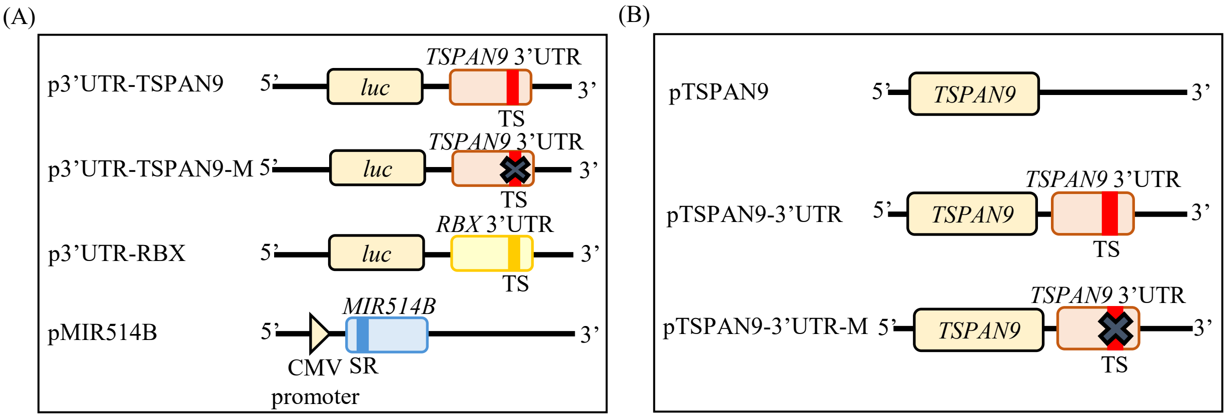


**Fig. S9. Details of different constructs used in the present study.** (A) Conservation of TS in the 3'UTR of *TSPAN9* across species. Note, the base complementarity of SR of miR-514b-3p with TS of *TSPAN9*. (B) A schematic representation of different constructs used in the dual-luciferase reporter assay. (C) A schematic representation of the TSPAN9 overexpression constructs. *Abbreviations*: TS, target site; and SR, seed region.

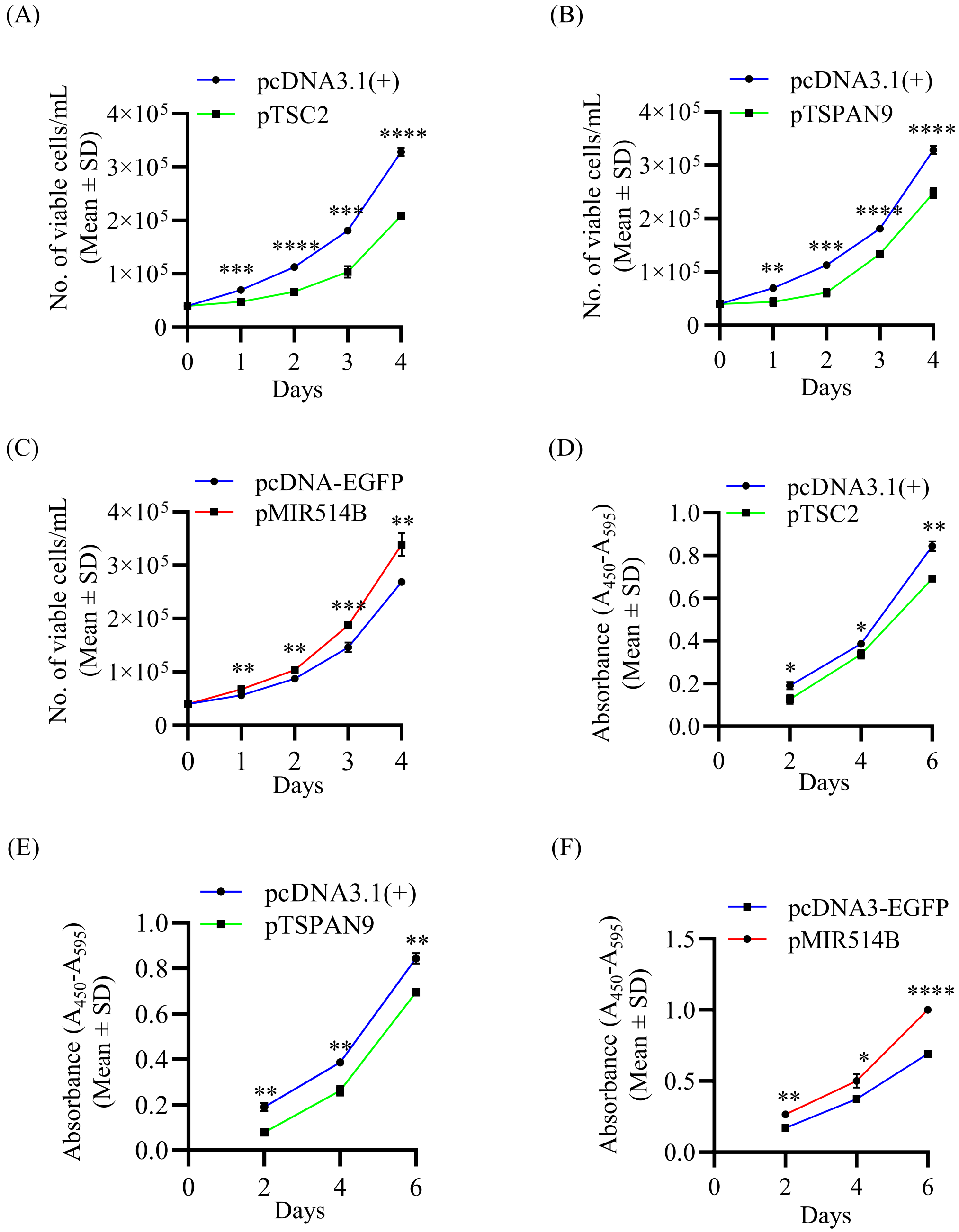


**Fig. S10. Effects of TSC2, TSPAN9 and miR-514b-3p on proliferation of SCC084 cells.** (A-C) The trypan blue dye exclusion assay showing the rate of proliferation of cells transfected with pTSC2, pTSPAN9 or pMIR514B (N=4). (D-F) The BrdU cell proliferation assay showing the rate of proliferation of cells transfected with pTSC2, pTSPAN9 or pMIR514B (N=3). P-values were calculated with Unpaired student's t-test with Welch correction.

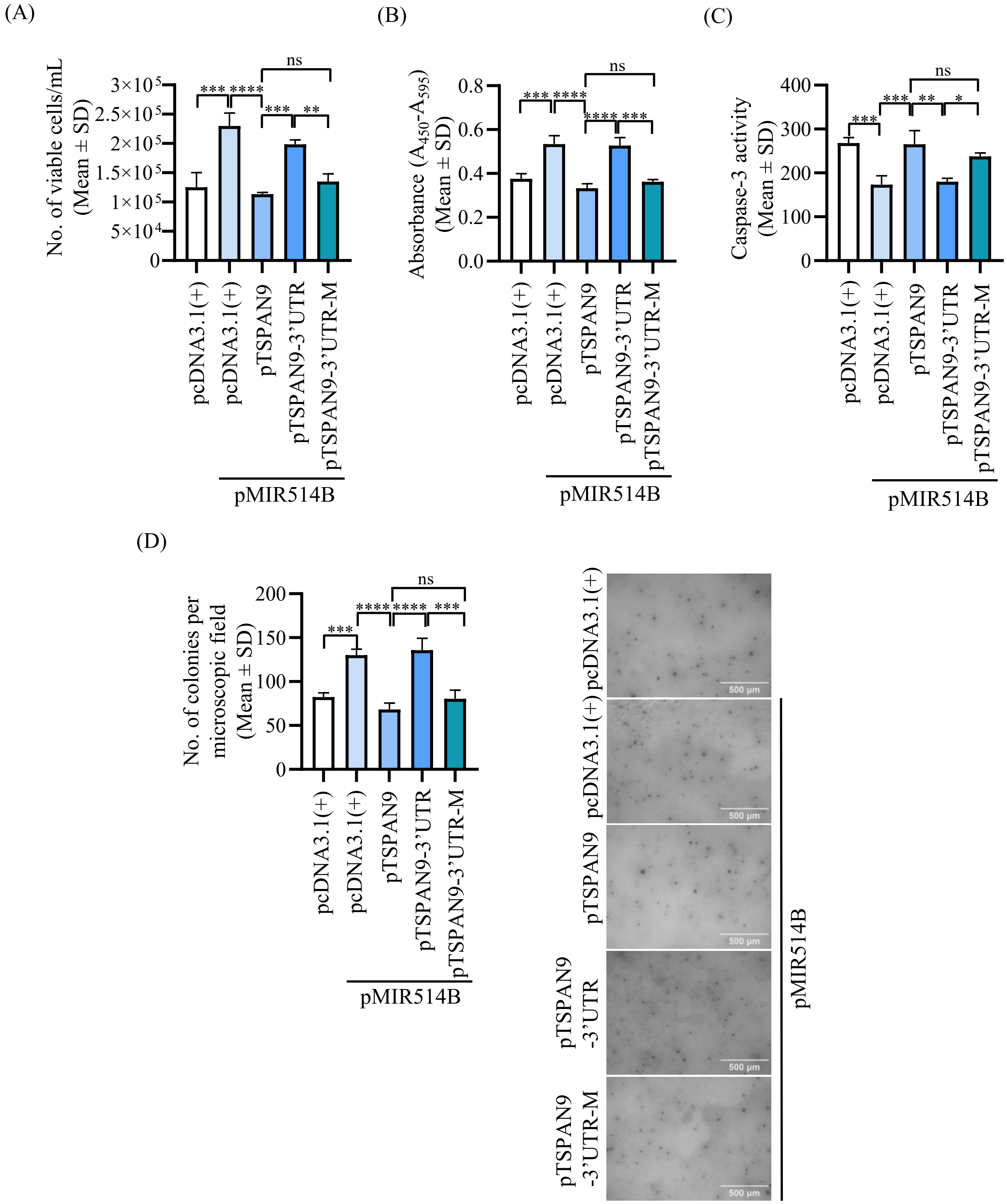


**Fig. S11. miR-514b-3p regulates proliferation, apoptosis, and anchorage-independent growth of SCC084 cells, in part, by targeting the 3'UTR of *TSPAN9*.** (A) Effects of miR-514b-3p and different *TSPAN9* constructs on proliferation of SCC131 cells by the Trypan blue dye exclusion assay (N=3) and (B) by the BrdU cell proliferation assay (N=3). (C) Effects of miR-514b-3p and different *TSPAN9* constructs on apoptosis by the Caspase-3 activation assay in SCC084 cells (N=4). (D) Effects of miR-514b-3p and different *TSPAN9* constructs on anchorage-independent growth of SCC084 cells by the soft agar colony formation assay (left panel). Microscopic images of the soft agar colony formation assay are also shown (right panel) (N=3). P-values were calculated with one way ANOVA with Tukey's multiple comparison test.

**Table S1: A list of differentially expressed miRNAs identified by the miRNA microarray analysis of TSC2 depleted SCC131 cells.**

**List of upregulated miRNAs**

| **Sl. no.** | **miRNA** | **Fold change expression** | **P-value** |
|---|---|---|---|
| 1. | hsa-miR-6880-5p | 52.61420757 | 0.000103 |
| **2.** | **hsa-miR-514b-3p** | **28.48987701** | **0.000004** |
| 3. | hsa-miR-181c-3p | 8.324497334 | 0.030125 |
| 4. | hsa-miR-885-3p | 4.695281694 | 0.088387 |
| 5. | hsa-miR-6825-5p | 4.452285598 | 0.069584 |
| 6. | hsa-miR-5187-5p | 3.669542522 | 0.156438 |
| 7. | hsa-miR-6509-5p | 3.166173037 | 0.009855 |
| 8. | hsa-miR-6873-3p | 2.515858101 | 0.000309 |
| 9. | hsa-miR-498 | 2.417963294 | 0.212724 |
| 10. | hsa-miR-3135b | 2.173806108 | 0.016253 |
| 11. | hsa-miR-6130 | 2.108666049 | 0.000462 |
| 12. | hsa-miR-4726-5p | 2.083111097 | 0.000005 |
| 13. | hsa-miR-6805-5p | 2.060477416 | 0.000111 |
| 14. | hsa-miR-3692-5p | 2.033441939 | 0.000491 |
| 15. | hsa-miR-6817-5p | 2.009385245 | 0.000065 |
| 16. | hsa-miR-6847-5p | 2.000693367 | 0.000193 |
| 17. | hsa-miR-205-3p | 1.972460667 | 0.000053 |
| 18. | hsa-miR-8071 | 1.889109859 | 0.000184 |
| 19. | hsa-miR-6849-5p | 1.8618627 | 0.000233 |

**List of downregulated miRNAs**

| Sl. no. | miRNA | Fold change expression | P-value |
|---|---|---|---|
| 1. | hsa-miR-3681-3p | 0.020677549 | 0.000181 |
| 2. | hsa-miR-4664-5p | 0.03010803 | 0.000000 |
| 3. | hsa-miR-6892-3p | 0.039618755 | 0.000002 |
| 4. | hsa-miR-550a-3-5p | 0.041390349 | 0.000001 |
| 5. | hsa-miR-6833-3p | 0.143118894 | 0.052502 |
| 6. | hsa-miR-8064 | 0.165133663 | 0.052851 |
| 7. | hsa-miR-758-5p | 0.176247377 | 0.065980 |
| 8. | hsa-miR-491-5p | 0.214796646 | 0.056582 |
| 9. | hsa-miR-6881-3p | 0.305163139 | 0.028909 |
| 10. | hsa-miR-6778-3p | 0.331279265 | 0.014497 |
| 11. | hsa-miR-663b | 0.33882021 | 0.001726 |
| 12. | hsa-miR-6514-3p | 0.419409004 | 0.000027 |
| 13. | hsa-miR-501-3p | 0.420488735 | 0.227657 |
| 14. | hsa-miR-1250-3p | 0.429284919 | 0.000643 |
| 15. | hsa-miR-34b-3p | 0.452700689 | 0.000637 |
| 16. | hsa-miR-5684 | 0.459653892 | 0.264568 |
| 17. | hsa-miR-451b | 0.473712111 | 0.000069 |
| 18 | hsa-miR-5010-3p | 0.487539967 | 0.000096 |
| 19. | hsa-miR-4446-5p | 0.489348971 | 0.000359 |
| 20. | hsa-miR-4730 | 0.493729431 | 0.000001 |
| 21. | hsa-miR-6815-3p | 0.50581388 | 0.161485 |
| 22. | hsa-miR-6716-3p | 0.518711692 | 0.000039 |
| 23. | hsa-miR-3591-3p | 0.540614701 | 0.000044 |
| 24. | hsa-miR-3200-3p | 0.560579877 | 0.020111 |

**Table S2: Details of primers used in RT-qPCR.**

| Sl. no. | Target Gene | Primer sequence (5' to 3') | Amplicon size (bp) | Annealing temp. (°C) |
|---|---|---|---|---|
| 1. | *TSC2* | Fw: CCGTCTTCCACATCGCCACCCTG | 168 | 60 |
| | | Rv: ACGATCACGTGGACAAAGTTGAACTG | | |
| 2. | *EREG** | Fw: ACAACTGTGATTCCATCATGTATCC | 109 | 62 |
| | | Rv: CTACACTTTGTTATTGACACTTGAGC | | |
| 3. | *GAPDH** | Fw: GAAGGGTGAAGGTCGGAGTC | 226 | 60 |
| | | Rv: GAAGATGGTGATGGGATTTC | | |
| 4. | miR-514b-3p | RT6-miR-514b-3p: TGTCAGGCAACC GTATTCACCGTGAGTGGTTCCACT | 84 | 56 |
| | | shortmiR-514b-3p: CGTCAGATGTCCG AGTAGAGGGGGAACGGCGATTGAC ACCTCTGTGAGT | | |
| | | MP Fw: TGTCAGGCAACCGTATTCACC | | |
| | | MP Rv: CGTCAGATGTCCGAGTAGAGG | | |
| 5. | *5S rRNA*** | Fw: GCCCGATCTCGTCTGATCT | 93 | 60 |
| | | Rv: AGCCTACAGCACCCGGTATT | | |
| 6. | *TSPAN9* | Fw: GTCATTGCCATAGGCACCATTGTC | 175 | 56 |
| | | Rv: CGTTCTCGTTCACCTTGTCCATG | | |
| 7. | *GULP1* | Fw: GCACCTCCAGCAGGCAGTATGAC | 173 | 58 |
| | | Rv: GGTTCTGCTCCAAACAGGTCCCG | | |
| 8. | *PTEN* | Fw: CCCAGTCAGAGGCGCTATGTG | 142 | 56 |
| | | Rv: CAAACTGAGGATTGCAAGTTCCG | | |
| 9. | *SESN3* | Fw: CAGAGAAGGAAGTTGTCCAAGC | 160 | 58 |
| | | Rv: CGCAACATGTAAAACTGGCTCC | | |
| 10. | *CABLES1* | Fw: GGACGGAGGAAGACAATCAACTGG | 151 | 58 |
| | | Rv: GTCTATGGTATTTCTCCGGGCTCC | | |
| 11. | *RBX* | Fw: CTGTGCCATCTGCAGGAACCAC | 142 | 56 |
| | | Rv: GAGCCAGCGAGAGATGCAGTGG | | |

*Abbreviations:* Fw, forward primer; Rv, reverse primer; bp, base pair; and temp, temperature.

**Table S3: Details of primers used in ChIP-qPCR.**

| Sl. no. | Target region | Primer sequence (5' to 3') | Amplicon size (bp) | Annealing temp ($^0C$) |
|---|---|---|---|---|
| 1. | *EREG* promoter | Fw: GTAAGTCACACGCACAGCTCTCC | 140 | 60 |
| | | Rv: CTTGGTTTGCTTTGTGGGGTCAC | | |
| 2. | TBS1 | Fw: CTATGGTAATCAGTACGCGATG | 137 | 58 |
| | | Rv: CAGCCCAGCCTAATGACTGAC | | |
| 3. | TBS2 | Fw: GACAGTGTAGTGCATGCTACAG | 142 | 58 |
| | | Rv: GATAGCCAGAGCTTATCTATTACG | | |
| 4. | TBS3 | Fw: CGTAATAGATAAGCTCTGGCTATC | 190 | 58 |
| | | Rv: CATCGCGTACTGATTACCATAG | | |
| 5. | TBS4 | Fw: CTCTCAAGCTTTGGGAAGGTGC | 146 | 58 |
| | | Rv: CCCCAGAATAATCATAAAACAGAG | | |
| 6. | TBS5 | Fw: CTGTGTTGTTACCTGGAGCATCG | 157 | 58 |
| | | Rv: GCTTGAGAGCAGTGAGTCAGAGA | | |
| 7. | TBS6 | Fw: CAAGTCTTTCAGGTTTGTTCTGAT | 167 | 58 |
| | | Rv: CGATGCTCCAGGTAACAACACAG | | |
| 8. | TBS7 | Fw: CTGTTCTTTAGAGAGCACAGCC | 128 | 56 |
| | | Rv: CTGTGTGATACACAGAGACCAC | | |

*Abbreviations:* Fw, forward primer; Rv, reverse primer; bp, base pair; and temp, temperature.

**Table S4:** ***In silico*** **prediction of gene targets of miR-514b-3p by target prediction tools.**

| Sl. No. | Agilent GeneSpring GX | miRDB | TargetScan Human8.0 | microT-CDS | STarMirDB |
|---|---|---|---|---|---|
| 1. | ***TSPAN9**** | ***TSPAN9**** | ***TSPAN9**** | ***TSPAN9**** | ***TSPAN9**** |
| 2. | **-** | ***GULP1**** | ***GULP1**** | ***GULP1**** | ***GULP1**** |
| 3. | **-** | ***CABLES1**** | ***CABLES1**** | **-** | ***CABLES1**** |
| 4. | **-** | ***PTEN**** | ***PTEN**** | ***PTEN**** | **-** |
| 5. | **-** | ***SESN3**** | ***SESN3**** | ***SESN3**** | ***SESN3**** |
| 6. | *NDUFA10* | *NCOA7* | *CDR1as* | *BAALC* | *ZNF831* |
| 7. | *C11orf49* | *ANO5* | *LPAR4* | *KIAA1549* | *NRXN3* |
| 8. | *ZNF491* | *PTPRG* | *DEFB132* | *TMEM260* | *STXBP5L* |
| 9. | *GALP* | *CD200R1* | *CD200R1* | *ZNF474* | *NR1D2* |
| 10. | *WDYHV1* | *TMEM68* | *AGBL3* | *AGO4* | *ARHGEF9* |
| 11. | *SPATS2L* | *JAM2* | *RBX1* | *PEG3* | *ZNF770* |
| 12. | *IDS* | *EML6* | *SPDYA* | *GRIA2* | *ZC3H14* |
| 13. | *RGPD3* | *EGFR* | *OR2L2* | *RBX1* | *ERMN* |
| 14. | *TMEM260* | *ERC1* | *TMEM50A* | *TSPAN12* | *DIP2B* |
| 15. | *SERF1B* | *RBX-1* | *CAPSL* | *ARHGEF28* | *SPTLC3* |

*The genes highlighted in bold are chosen as putative gene targets of miR-514b-3p for experimental validation.

***Abbreviations*****:** *NDUFA10*, NADH Ubiquinone Oxidoreductase Subunit A10; *TSPAN9*, Tetraspanin 9; *C11orf49*, Chromosome 11 Open Reading Frame 49; *GULP1*, GULP PTB Domain Containing Engulfment Adaptor 1; *CABLES1*, Cdk5 and Abl Enzyme Substrate 1; *ZNF491*, Zinc Finger Protein 491; *PTEN*, Phosphatase and Tensin Homolog; *GALP*, Galanin Like Peptide; *SESN3*, Sestrin 3; *WDYHV1*, WDYHV Motif Containing 1; *NCOA7*, Nuclear Receptor Coactivator 7; *CDR1as*, Cerebellar Degeneration-Related Protein 1 Antisense; *BAALC*, BAALC Binder Of MAP3K1 And KLF4; *ZNF831*, Zinc Finger Protein 831; *SPATS2L*, Speriolin Like Spermatogenesis Associated 2; *ANO5*, Anoctamin 5; *LPAR4*, Lysophosphatidic Acid Receptor 4; *KIAA1549*, *KIAA1549* Gene; *NRXN3*, Neurexin 3; *IDS*, Iduronate 2-Sulfatase; *PTPRG*, Protein Tyrosine Phosphatase Receptor Type G; *DEFB132*, Defensin Beta 132; *TMEM260*, Transmembrane Protein 260; *STXBP5L*, Syntaxin Binding Protein 5 Like; *RGPD3*, RANBP2-Like And GRIP Domain Containing 3; *CD200R1*, CD200 Receptor 1; *ZNF474*, Zinc Finger Protein 474; *NR1D2*, Nuclear Receptor Subfamily 1 Group D Member 2; *TMEM68*, Transmembrane Protein 68; *AGBL3*, ATP/GTP Binding Protein Like 3; *AGO4*, Argonaute RISC Catalytic Component 4; *ARHGEF9*, Rho Guanine Nucleotide Exchange Factor 9; *SERF1B*, Small EDRK-Rich Factor 1B; *JAM2*, Junctional Adhesion Molecule 2; *RBX1*, Ring-Box 1, E3 Ubiquitin Ligase; *PEG3*, Paternally Expressed 3;

*ZNF770*, Zinc Finger Protein 770; *SERF1A*, Small EDRK-Rich Factor 1A; *EML6*, Echinoderm Microtubule Associated Protein Like 6; *SPDYA*, Speedy/RINGO Cell Cycle Regulator Family Member A; *GRIA2*, Glutamate Ionotropic Receptor AMPA Type Subunit 2; *ZC3H14*, Zinc Finger CCCH-Type Containing 14; *POU2F1*, POU Class 2 Homeobox 1; *EGFR*, Epidermal Growth Factor Receptor; *OR2L2*, Olfactory Receptor Family 2 Subfamily L Member 2; *RGPD4*, RANBP2-Like And GRIP Domain Containing 4; *ERC1*, ELKS/Rab6-Interacting/CAST Family Member 1; *TMEM50A*, Transmembrane Protein 50A; *TSPAN12*, Tetraspanin 12; *DIP2B*, Disco-Interacting Protein 2B; *GCC2*, GRIP And Coiled-Coil Domain Containing 2; *CAPSL*, Calcyphosine Like; *ARHGEF28*, Rho Guanine Nucleotide Exchange Factor 28; and, *SPTLC3*, Serine Palmitoyl transferase Long Chain Base Subunit 3.

**Table S5: Details of plasmid constructs generated in the present study.**

| Sl. no. | Construct | Cloning vector | Primer sequence (5' to 3') | Amplicon size (bp) | Annealing temp. (°C) |
|---|---|---|---|---|---|
| 1. | pPROM-MIR514B | pGL3-basic | Fw: GCTAGGTACCCCCTCTCATGGGCATTTTCTGTC<br>*Kpn* I<br>Rv: TAGCACGCGTCTTTCAGTTTCTAGGTCTTACGC<br>*Mlu* I | 2,825 | 56 |
| 2. | pMIR514B | pcDNA3-EGFP | Fw: GCTAAAGCTTCTATGGTAATCAGTACGCGATG<br>*Hind* III<br>Rv: TAGCCTCGAGGAACAGGGAGGGTGTCAGTAG<br>*Xho* I | 350 | 56 |
| 3. | p3'UTR-TSPAN9 | pMIR-REPORT | Fw: GCTAAAGCTTGTCTGGGTGGATGGAAGTGGCTG<br>*Hind* III<br>Rv: TAGCGAGCTCGGTGTGTCTTGGATGAAGGGTAGG<br>*Sac* I | 591 | 54 |
| 4. | p3'UTR-RBX | pMIR-REPORT | Fw: GCTAACGCGTGTATGGGCACTAGGAAAAGACTTC<br>*Mlu* I<br>Rv: TAGCGAGCTCCAGAAGCCCACAGTACAGAGGAG<br>*Sac* I | 563 | 54 |
| 5. | pTSPAN9 | pcDNA 3.1(+) | Fw: GCTAAAGCTTATGGCCAGGGGCTGCCTCTGCTG<br>*Hind* III<br>Rv: TAGCGAATTCTCATGCGTCGTACTTCTTACCAGTC<br>*Eco* RI | 720 | 65 |
| 6. | pTSPAN9-3'UTR | pTSPAN9 | Fw: GCTAGAATTCGTCTGGGTGGATGGAAGTGGCTG<br>*Eco* RI<br>Rv: TAGCGCGGCCGCGGTGTGTCTTGGATGAAGGGTAGG<br>*Not* I | 591 | 56 |
| 7. | pTSPAN9-3'UTR-M | pTSPAN9 | Fw: GCTAGAATTCGTCTGGGTGGATGGAAGTGGCTG<br>*Eco* RI<br>Rv: TAGCGCGGCCGCGGTGTGTCTTGGATGAAGGGTAGG<br>*Not* I | 583 | 56 |
| **Primers used in the site-directed mutagenesis** | | | | | |
| 1. | p3'UTR-TSPAN9-M | pMIR-REPORT | Fw: ATAAAAACGGATAAAAGCACAGCAGGCTGT<br>Rv: ACAGCCTGCTGTGCTTTTATCCGTTTTTAT | 583 | 56 |

*Abbreviations:* Fw, forward primer; Rv, reverse primer; bp, base pair; and temp, temperature.

**Table S6: Details of Mock and TSC2 siRNAs used for TSC2 knockdown.**

| Sl. no. | siRNA name | siRNA sequence (5’ to 3’) |
|---|---|---|
| 1. | ON-TARGET plus$^{TM}$ Non-Targeting Pool (Mock siRNAs) Cat# D-001810-10-05 | TGGTTTACATGTCGACTAA<br>TGGTTTACATGTTGTGTGA<br>TGGTTTACATGTTTTCTGA<br>TGGTTTACATGTTTTCCTA |
| 2. | ON-TARGET plus$^{TM}$ Human TSC2 (7249) siRNA-SMART Pool Cat# L-003029-00-0005 | GGAATGTGGCCTCAACAAT<br>GGATTACCCTTCCAACGAA<br>CGAACGAGGTGGTGTCCTA<br>GCATTAATCTCTTACCATA |

**Table S7: Details of the antibodies used in the present study.**

| Primary antibody | | | | | |
|---|---|---|---|---|---|
| **Sl. no.** | **Antibody** | **Dilution used** | **Observed band size** | **Catalog #** | **Manufacturer** |
| 1. | Anti-TSC2 (Rabbit) | 1:10,000 | 200 kDa | ab52936 | Abcam, Cambridge, UK |
| 2. | Anti-TSC1 (Rabbit) | 1:1000 | 130 kDa | 6935S | Cell Signaling Technology, Danvers, USA |
| 3. | Anti-Phospho TSC2 Ser 939 (Rabbit) | 1:1,000 | 200 kDa | ab52962 | Abcam, Cambridge, UK |
| 4. | Anti-TSPAN9 (Rabbit) | 1:10,000 | 35 kDa | PA5-20874 | Thermo Fisher Scientific, Waltham, MA, USA |
| 5. | Anti-β actin (Mouse) | 1:10,000 | 42 kDa | A5441 | Sigma-Aldrich, St. Louis, MO, USA |
| 6. | Anti-Total S6K1 (Rabbit) | 1:1,000 | 70 kDa | 9202S | Cell Signaling Technology, Danvers, USA |
| 7. | Anti-Phospho S6K1 Thr 389 (Rabbit) | 1:1,000 | 70 kDa | 9205S; 9234S | Cell Signaling Technology, Danvers, USA |
| 8. | Anti-Total AKT (Rabbit) | 1:1,000 | 60 kDa | 4685S | Cell Signaling Technology, Danvers, USA |
| 9. | Anti-Phospho AKT Ser 473 (Rabbit) | 1:2,000 | 60 kDa | 4060S | Cell Signaling Technology, Danvers, USA |
| 10. | Anti-p55 (Rabbit) | 1:1,000 | 55 kDa | 11889S | Cell Signaling Technology, Danvers, USA |
| 11. | Anti-H3 (Rabbit) | 1:10,000 | 17 kDa | 4620S | Cell Signaling Technology, Danvers, USA |
| 12. | Anti-GAPDH (Mouse) | 1:25,000 | 37 kDa | G8785 | Sigma-Aldrich, St. Louis, MO, USA |
| **Secondary antibody** | | | | | |
| 1. | Anti-mouse IgG HRP-conjugated | 1:2,500 | 105502 | | Bangalore Genei, Bangalore, India |
| 2. | Anti-rabbit IgG HRP-conjugated | 1:2,500 | 105499 | | Bangalore Genei, Bangalore, India |

*Abbreviation:* kDa, kilo Dalton.
